# Demethylating agents drive PARP inhibitor resistance in ovarian carcinomas with *BRCA1* gene silencing

**DOI:** 10.1101/2025.04.07.647418

**Authors:** Ksenija Nesic, Franziska Geissler, Lijun Xu, Elizabeth Kyran, Sally Beard, Cassandra J. Vandenberg, Brett Liddell, Sam W.Z. Olechnowicz, Monique Topp, Orla McNally, Gayanie Ratnayake, Nadia Traficante, Australian Ovarian Cancer Study, Anna DeFazio, David D. L. Bowtell, Tony Pappenfuss, Fan Zhang, Alexander Dobrovic, Nicola Waddell, Clare L. Scott, Olga Kondrashova, Matthew J. Wakefield

## Abstract

DNA methyltransferase 1 inhibitor (DNMT1i) therapy is a promising option for increasing immune response as part of combination cancer therapy. High-grade serous ovarian carcinoma (HGSOC) is a highly aggressive cancer with poor survival outcomes, where DNMT1i therapy is being increasingly explored. HGSOC with epigenetically silenced *BRCA1* has been shown to respond to PARP inhibitor (PARPi) treatment – a core targeted therapy for HGSOC. However, loss of silencing of even a single *BRCA1* allele causes PARPi and platinum chemotherapy resistance. We tested whether *BRCA1* silencing was robust to DNMT1i therapy, or would be reversed, thus driving PARPi resistance.

We previously generated two homozygously silenced *BRCA1* HGSOC cell lines: WEHI-CS62 and an OVCAR8 derivative. DNMT1i treatment caused sustained *BRCA1* promoter methylation loss, gene re-expression and PARPi resistance in both of these silenced *BRCA1* lines, but not in mutated *BRCA1/2* or *RAD51C* lines. Methylation arrays confirmed transient global CpG methylation losses following DNMT1i. CRISPR deletion of the re-expressed *BRCA1* copy in WEHI-CS62 restored silencing and PARPi sensitivity. Furthermore, DNMT1i treatment of a silenced *BRCA1* PDX caused heterogeneous *BRCA1* promoter methylation loss.

In summary, DNMT1 inhibitors caused sustained reduction of *BRCA1* promoter methylation in HGSOC cells. This resulted in *BRCA1* re-expression and PARP inhibitor resistance, presenting a significant risk for up to 17% of HGSOC patients with *BRCA1* gene silencing who could benefit from PARP inhibitor therapy. We conclude that DNA demethylation therapy should be avoided for HGSOC patients with epigenetically silenced *BRCA1*.

## Introduction

PARPi, a type of DNA damage repair inhibitor (DDRi), specifically targets poly(ADP-ribose) polymerase (PARP) enzymes and has emerged as a breakthrough therapeutic option for cancers with DNA double-strand break repair deficiencies. While the synthetic lethality of PARPi with specific genetic alterations, such as *BRCA1* and *BRCA2* mutations, has dramatically improved outcomes for patients, combination therapies will be required to increase the number of patients that achieve durable responses.

One combination therapy approach that is being increasingly explored in solid cancers, including ovarian cancer, is the inhibition of DNA methyltransferase 1 (DNMT1i) [1, 2]. Clinical trials in ovarian cancer include NCT00529022, NCT00477386, and NCT02159820, and those in advanced solid cancers include NCT02811497, NCT05320640, and NCT02998567. DNMT1i has been shown to improve immune checkpoint blockade-based cancer therapy via a diverse array of mechanisms, including unmasking of silenced endogenous retroviruses (ERVs), and expression of chemokine Th1 or MHC, thereby enabling neoantigen presentation to antigen-presenting cells [1, 2]. Conditional loss of DNMT1 has also been shown to alter the tumour microenvironment to allow T cell infiltration, thus reducing tumour growth and enhancing immune checkpoint blockade [3]. One hypothesis supporting the use of DNMT1i in ovarian cancer is their potential effect on immune priming of cancer cells and their microenvironment [4–7], with several clinical trials completed or underway (e.g. NCT02901899, NCT01696032, NCT03206047). Another is their ability to potentially reverse platinum resistance by reactivating genes that have been silenced as a mechanism of acquired drug resistance, with some promising evidence in early clinical trials [8–10]. Combination DNMT1i and PARPi treatment in breast and ovarian cell lines and xenograft models has also been shown to enhance cytotoxicity independent of *BRCA* mutation status, by increasing PARP trapping at DNA double strand break sites [11].

5-azacytidine (5-AZA) and 5-aza-2’deoxycytidine (decitabine) are DNA demethylating agents that inhibit DNA methyltransferase 1 (DNMT1) by trapping it on DNA, leading to impaired methylation maintenance during DNA replication, and subsequent global DNA hypomethylation [2]. New nucleoside DNMT1i with improved stability and bioavailability (*e.g.* guadecitabine), as well as non-nucleoside DNMT1i, have potential to significantly enhance the utility of this drug class. GSK3484862 and GSK3685032 are potent, selective, small molecule DNMT1i with low cellular toxicity, which have recently demonstrated potent demethylation capacity in pre-clinical *in vitro* and *in vivo* models [12–14]. In mice, GSK3685032 demonstrated improved tolerability and increased DNA hypomethylation compared to decitabine, resulting in enhanced anti-tumour activity in murine models of AML [14].

In HGSOC epigenetic silencing of *BRCA1* is observed in up to 17% of cases [15]. This frequency is only exceeded as a cause of homologous recombination deficiency (HRD) by *BRCA1/2* mutations, which are observed in ∼ 20% of cases [16, 17]. Epigenetic silencing is characterized by homozygous *BRCA1* promoter methylation (me*BRCA1*), and correlates with response to PARPi [15]. Partial loss of me*BRCA1* following platinum chemotherapy or PARPi indicates gene re-expression and is a proven mechanism of loss of HRD and acquired PARPi resistance in the clinic [15, 18, 19]. However, it is not known what mechanisms lead to loss of silencing of *BRCA1* following platinum chemotherapy or PARPi. Indeed, some patients with silenced *BRCA1* can demonstrate very long-term and durable responses to PARPi, while others lose methylation and re-express *BRCA1* relatively rapidly [15, 18–20]. It is possible that PARPi and/or platinum directly cause loss of *BRCA1* silencing *-* generating new drug-resistant HR-proficient (HRP) clones - and provide strong selection of new or pre-existing HRP drug-resistant clones.

We hypothesised that silencing of *BRCA1* may involve mechanisms beyond methylation and be robust to treatment with DNMT1i. Alternatively, treatment with DNMT1i may pose a risk to patients with silenced *BRCA1* by causing demethylation of DNA, gene re-expression and loss of HRD - thus rendering their cancers HRP and profoundly PARPi-resistant.

To test the stability of silencing under the pressure of DNMT1i treatment, we used two HGSOC cell lines with homozygous me*BRCA1* resulting in gene silencing, originating from our laboratory. In both homozygously silenced *BRCA1* and PARPi-responsive cell lines, treatment with DNMT1i led to heterogeneous *BRCA1* methylation loss, *BRCA1* gene re-expression, and resistance to PARPi. PARPi resistance was not observed in three cell lines with HRD without homozygous me*BRCA1* treated with DNMT1i. Knock-out of unmethylated *BRCA1* epialleles in a post-DNMT1i cell line confirmed that *BRCA1* re-expression drove resistance. DNMT1i treatment also caused heterogeneous reduction of me*BRCA1 in vivo* in PDX.

We show DNMT1i could reverse silencing of *BRCA1* in HGSOC cell lines and PDX containing homozygous promoter methylation and silencing of *BRCA1*. This includes a cell line with highly stable *BRCA1* silencing, WEHI-CS62. Even partial loss of *BRCA1* methylation led to re-expression of *BRCA1* and PARPi resistance. Thus, DNA demethylating therapy should be avoided in HGSOC patients with me*BRCA1* or HRD with an unknown driver, in order to preserve the therapeutically targetable HRD and PARPi response.

## Methods

### Reagents and Cell lines

Rucaparib was kindly provided by Clovis Oncology (USA). GSK3484862 was purchased from MedChemExpress (Catalog # HY-135146) and GSK3685032 was purchased from TargetMol (Catalog # T9573). Propidium iodide (PI) for FACS was purchased from Sigma-Aldrich (Catalog # P4864-10ML). The parental OVCAR8 cell line was kindly gifted to us from Clovis Oncology in 2016, and all OVCAR8 variants described in this study were generated by our laboratory [21, 22]. The PEO1 cell line was kindly gifted to us from Scott Kaufmann (Mayo Clinic, MN, USA). The WEHI-CS62 cell line was generated in our laboratory from a PDX tumour, as previously described [15]. Cell line identities were confirmed using STR profiling (AGRF). Cell lines were confirmed to be negative for Mycoplasma during culture using the MycoAlert Mycoplasma Detection Kit (Lonza, Cat# LT07-118).

### DNMTi treatment of cell lines

Cells were seeded on 6-well plates or T25 flasks and treated with indicated doses of GSK3484862, GSK3685032 or decitabine for 4-5 days (as indicated), with media/drug replaced every day during this period. The DMSO control concentration was equivalent to the volume of drug (made in DMSO) used at the highest concentration.

### Nucleic acid extractions

The Qiagen AllPrep Mini Kit (Catalog # 80204) was used to extract DNA and RNA for all experiments on 6-well plates. For cells corresponding to Incucyte drug response experiments, plate based extraction methods from Zymo Research were used to extract DNA (Catalog # D4070) and RNA (Catalog # R1052) according to manufacturer’s protocols.

### Methylation-specific droplet digital PCR and high resolution melt analysis

Methylation-specific droplet digital PCR and high-resolution melt analysis for *BRCA1* promoter methylation levels was performed as previously described in Kondrashova et al., 2018 [15].

### Targeted *BRCA1* bisulfite sequencing

Targeted bisulfite sequencing of the *BRCA1* promoter was performed as previously described [23] using a two-step PCR protocol. New primers specific to the *BRCA1* promoter region hg38 chr17:43125457 – 43125336 were designed for this assay (based on primers from [24], with Illumina sequencing adaptors added):

Forward primer (5’-3’), Illumina sequencing adaptors as bolded text: **TCGTCGGCAGCGTCAGATGTGTATAAGAGACAG**TtgTtgTttagcggtagTTTTttggt; Reverse primer (5’-3’), Illumina sequencing adaptors as bolded text: **GTCTCGTGGGCTCGGAGATGTGTATAAGAGACAG**aAcctAtcccccgtccaAAaa

MeNGS data was analysed using the MethAmplicons package, using a 0.01 read frequency cut-off [23]. All samples were found to have an acceptably high bisulfite conversion efficiency (>98%), which reduces false positive methylation calls. “Fully methylated epialleles” were defined as epialleles with 8/9 CpG sites methylated for the *BRCA1* promoter.

### Reverse transcription quantitative real-time PCR

Reverse transcription quantitative real-time PCR for *BRCA1* gene expression was performed as described in Kondrashova et al., 2018 [15]. Briefly, RNA was converted to cDNA using Superscript III Reverse Transcriptase (Invitrogen), and qPCR was performed using SYBR Green PCR Master Mix (Applied Biosystems) following manufacturer’s instructions. Ct values for each sample were normalized to the average Ct values of four different housekeeping genes (*HPRT1, ACTB, SDHA*, and *GAPDH*), and resulting values were used to calculate fold-change of BRCA1 expression for each sample.

### Incucyte cell growth assays

Cells were plated in duplicate wells on Griener 384 well plates, with densities depending on the cell growth rate of each cell line (Summary provided in Supplementary Table S3). The day following seeding, cells were treated with indicated doses of rucaparib or AZD5305 and place on the Sartorius Incucyte S3 Live-Cell Analysis System where wells were imaged once every 12 hours for up to 14 days. For growth analysis, a cell confluence mask was applied to all wells at each timepoint and plotted using the built-in Incucyte software.

### CellTiter-Glo cell viability assays

Cells were plated in duplicate wells on Griener 384 well plates, with densities depending on the cell growth rate of each cell line (Summary provided in Supplementary Table S3). The day following seeding, cells were treated with indicated doses of AZD5305. After 16 days, 25 μl (1/4 final volume) of CellTiter-Glo 2.0 reagent (Promega) was added to cells in media, and placed on orbital shaker for 2 minutes. Plates were incubated at room temperature for 10 minutes before luminescence readings being taken on the CLARIOstar Plate Reader (BMG Labtech).

### EPIC methylation arrays and analysis

Detailed methods are available in the Supplementary Methods file.

### Fluorescence-Activated Cell Sorting

Cells were trypsinized, and resuspended in FACS buffer (DPBS with 7% fetal calf serum), then sorted as single cells into 96 well plates using propidium iodide to exclude dead cells. For post-GSK WEHI-CS62 clones, cells were treated with 5uM GSK for 5 days and left to recover over the weekend (2 days) prior to FACS.

### Ethical Approval

All experiments involving animals were performed according to the Code for the Care and Use of Animals for Scientific Purposes 8th Edition, 2013 (updated 2021), and were approved by the WEHI Animal Ethics Committee (2019.024 and 2022.030). Ovarian carcinoma PDX were generated from patients enrolled in the Australian Ovarian Cancer Study (AOCS)/WEHI Stafford Fox Rare Cancer Program (SFRCP). Informed written consent was obtained from all patients, and all experiments were performed according to human ethics guidelines. Additional ethics approval was obtained from the Human Research Ethics Committees at the Royal Women’s Hospital, the WEHI (HREC #10/05 and #G16/02) and QIMR Berghofer (P3456 and P2095).

### Establishment and treatment of PDX models

WEHI PDX #62 was established by transplanting viably frozen minced tumour subcutaneously into NOD.Cg*-Prkdc^scid^ Il2rg^tm1Wjl^/*SzJ (NSG; colony derived from Jackson Labs in 2018) recipient mice (T1, passage 1) as described previously [25]. Recipient mice bearing T2-T10 (passage 2 to passage 10) tumours were randomly assigned to treatments when tumour volume reached 180-300 mm^3^. *In vivo* cisplatin treatments were administered on days 1, 8 and 18, as previously described [26]. The regimen for rucaparib treatment was oral gavage once daily Monday-Friday for 3 weeks at 450 mg/kg for all models, as previously described [23]. The regimen for the DNMT1i GSK3685032 was sub-cutaneous injection (10µl/g) once daily on Monday, Wednesday & Friday for 4 weeks at 45mg/kg, as described previously [14]. Tumours were harvested once tumour volume reached 700 mm^3^ or when mice reached ethical or end of experiment (120 days post treatment) endpoints. Nadir, time to progression (TTP or PD), time to harvest (TTH), and treatment responses are as defined previously [15]. Data was plotted using the SurvivalVolume package [27].

### WEHI-CS62 Brunello CRISPR screens

Detailed methods are available in the Supplementary Methods file.

### Statistical analyses

An unpaired t test (two-tailed) was used to analyse ddPCR and RT-qPCR results (*P* values presented in supplementary Table S1 and S2) using PRISM (Graphpad). *P* < 0.05 was considered statistically significant and statistical tests are indicated in the figure legends. *P* values for PDX model treatments compared to vehicle treatment group were calculated by a log-rank test between fitted Kaplan-Meier estimates with a chi-squared null distribution and one degree of freedom using SurvivalVolumes v1.2.4 [27]. Two-tailed Wilcoxon signed-rank test were used to compare distribution of methylation from EPIC array.

## Results

### Silencing of *BRCA1* was robust to a genome-wide PARPi resistance CRISPR screen

To determine which pathways are critical to stability of *BRCA1* silencing we conducted a genome-wide PARPi resistance CRISPR screen in the WEHI-CS62 HGSOC cell line harbouring homozygous promoter methylation of *BRCA1* (me*BRCA1),* using the Brunello CRISPR guide library. During eight weeks of treatment with the PARPi, rucaparib, me*BRCA1* was retained in all screen replicates, consistent with high stability of this epigenetic mark in this cell line (Figure 1A-B). *DNMT1* is a core DNA methylation maintenance gene. No *DNMT1* guides were detected at the end of the screen in either PARPi treated or control samples (Figure 1C; Supplementary Figure S1A), suggesting that loss of *DNMT1* negatively impacted cell survival over time. As *DNMT1* loss is reportedly lethal to somatic dividing cells [28] we analysed DepMap data demonstrating a high level of essentiality for *DNMT1* in cancer cell lines (Figure 1D) and assessed the impact of *DNMT1* gene expression on patient survival using the TCGA ovarian cancer dataset (in GEPIA), which revealed that high *DNMT1* was associated with improved survival (p= 0.023; Supplementary Figure S1B), suggesting inhibition may be deleterious. We further investigated the effect of DNMT1 inhibition in HGSOC cell lines, with and without me*BRCA1,* to determine whether transient DNMT1 inhibition would lead to me*BRCA1* loss and PARPi resistance. This includes WEHI-CS62, which we determined via this CRISPR screen, has extremely stable me*BRCA1*.

**Figure 1.**
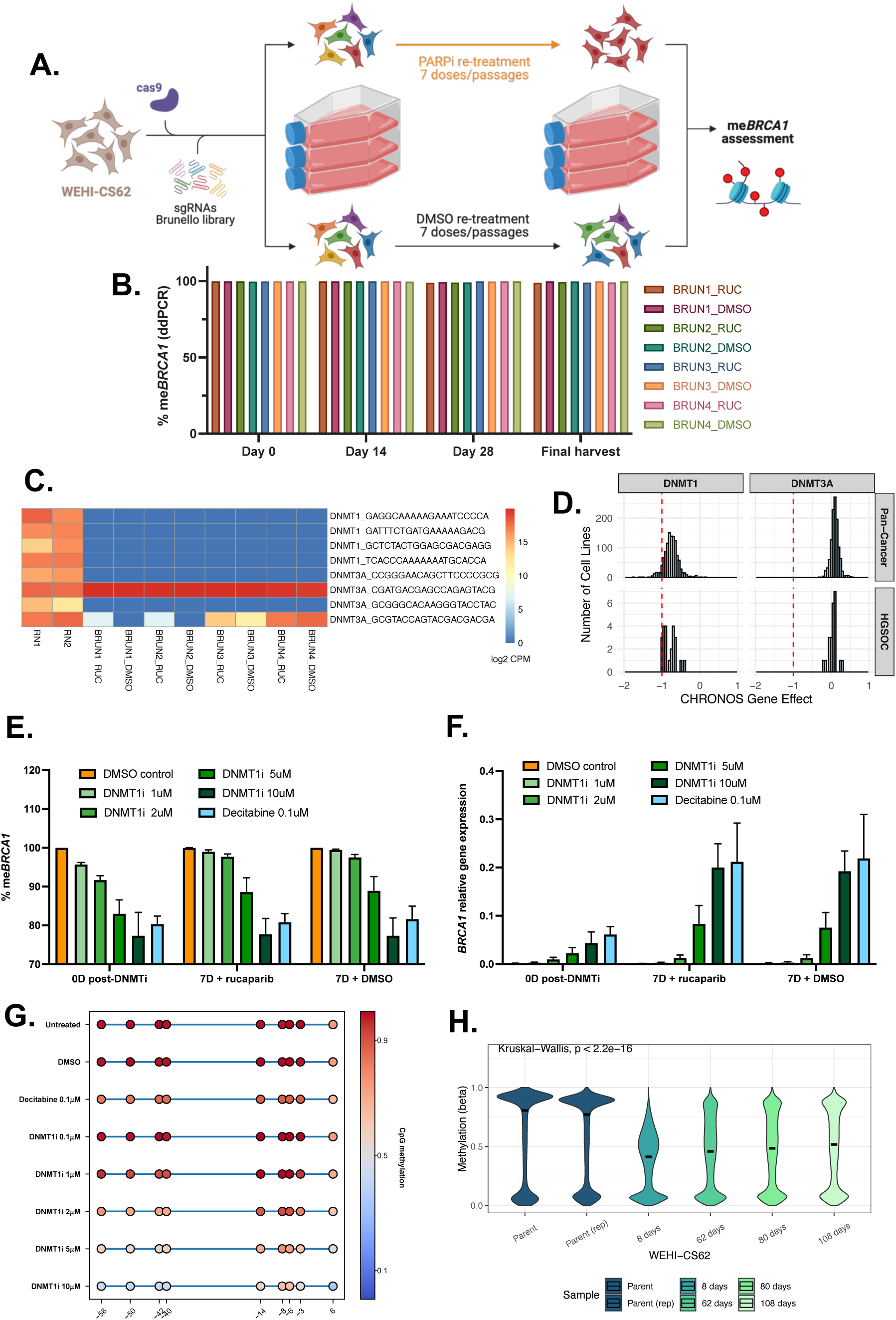
Transient DNMT1 inhibition caused *BRCA1* methylation loss and restored *BRCA1* gene expression in HGSOC cell line WEHI-CS62. (A) A genome-wide PARPi resistance CRISPR screen (Brunello guide library) in the homozygously methylated-*BRCA1* (me*BRCA1*) WEHI-CS62 cell line. Cells expressing Cas9 and the Brunello single guide RNA (sgRNA) library were exposed to seven doses of PARPi (rucaparib; RUC) or control (DMSO) for up to eight weeks. Cells were collected at days 0, 14, 28 and final harvest (56d for rucaparib-treated cells and 49d for DMSO control cells). Figure created using BioRender.com. (B) No significant me*BRCA1* loss (>1%) was observed in any replicates (BRUN1 to BRUN4) or treatment conditions at any timepoints using methylation-specific digital droplet PCR (MS-ddPCR). (C) *DNMT1* guide counts were depleted in treated and untreated final CRISPR screen samples, unlike DNMT3A (RN1 and RN2 represent pre-PARPi infection replicate 1 and 2 library diversity checks). (D) Gene essentiality analysis of *DNMT1* and *DNMT3A* (DepMap Public 24Q2 dataset). Each score represents the dependency of cell lines on the given gene (−1 scores, dashed line, indicate essentiality) using the Chronos computational method to analyse data from multiple CRISPR screens. Data for both pan-cancer (n= 1150) and HGSOC (n= 23) cell lines is presented. (E) me*BRCA1* was lost in WEHI-CS62 when cells were treated for four days with the DNA demethylating agent, decitabine, or with a targeted DNMT1 inhibitor (DNMT1i; % methylated epialleles measured by MS-ddPCR). Mean of n=3 experiments, with error bars showing standard deviation. P values presented in Supplementary Table S1. (F) increased *BRCA1* expression performed by reverse transcriptase qPCR of *BRCA1*. Mean of n=3 experiments, with error bars showing standard deviation. P values presented in Supplementary Table S1. (G) Patterns of me*BRCA1* loss across promoter CpG sites was assessed using targeted bisulfite next generation sequencing (meNGS) of the *BRCA1* promoter. Each circle represents average methylation level for a CpG site along the *BRCA1* promoter, and numbers on x-axis indicate CpG distance from *BRCA1* transcription start site. (H) Global DNA methylation losses in WEHI-CS62 following five days of GSK3484862 treatment were confirmed using Illumina EPIC global methylation arrays at indicated timepoints post-treatment. Parental (rep) is a replicate of WEHI-CS62 parent line of a different passage. The P value represents the significance in global methylation distribution difference across all samples using the Kruskal-Wallis test.

### DNMTi drove *BRCA1* promoter methylation loss and increased *BRCA1* expression in me*BRCA1* HGSOC

The WEHI-CS62 cell line, with two methylated gene copies of *BRCA1* (homozygous *BRCA1* [15]), was treated with a range of DNMT1i doses (GSK3484862 1-10 μM), 0.1 μM decitabine (DNMT-trapping nucleoside analogue; Supplementary Figure S2) or DMSO control for four days. The treated cells were either harvested immediately (0D post DNMTi) or subsequently treated with 1 μM PARPi rucaparib or DMSO control and left to recover for a further seven days prior to harvest (7D + rucaparib; 7D + DMSO). Cells treated with either DNMT1i or 0.1 μM decitabine for four days were found to have reduced me*BRCA1* by methylation-specific droplet digital PCR (MS-ddPCR; Figure 1E), and increased *BRCA1* expression by RT-qPCR (excluding DNMTi 1-2 μM doses Figure 1F; Supplementary Table S1). While me*BRCA1* levels did not decrease further at the later 7-day post treatment timepoints, *BRCA1* gene expression was significantly increased when compared to 0-days post-treatment (p values all <0.05, presented in Supplementary Table S1).

*BRCA1* promoter methylation levels at individual CpG sites and epiallele diversity following DNMT1i treatment in WEHI-CS62 was assessed using targeted bisulfite amplicon sequencing. Methylation across the CpG sites was heterogeneous following five days of decitabine or DNMT1i treatment, with me*BRCA1* levels inversely correlated with DNMT1i dose (Figure 1G). Fully unmethylated epialleles were present in DNMT1i 10 μM and decitabine 0.1 μM treated samples, and not observed in the DMSO control (Figure 1G; Supplementary Figure S3A-C). Global decreases in DNA methylation occurred in WEHI-CS62 eight days after DNMT1i treatment, (Figure 1H), and were not restricted to any particular CpG contexts (Supplementary Figure S3D). Of note, DNA methylation levels were found to recover over time for many CpG sites over 108 days (Figure 1H), but not for *BRCA1* (further EPIC array analysis presented in subsequent sections).

Human HGSOC cell line OVCAR8, which has heterozygous me*BRCA1* and *BRCA1* gene expression, was also treated with 10 μM DNMT1i and 0.1 μM decitabine, and had reduced methylation of the *BRCA1* promoter compared to DMSO control (Supplementary Figure S4; P values in Supplementary Table S2). There was also a trend towards increased *BRCA1* expression in DNMT1i or decitabine-treated OVCAR8 samples (Not significant; Supplementary Figure S4).

### DNMT1i treatment caused long-term instability of me*BRCA1*

Parental WEHI-CS62 was treated with DNMT1i (5 μM GSK3484862) for five days, allowed two days recovery and then sorted to generate single cell clones with distinct me*BRCA1* patterns that could be studied in the context of PARPi resistance (Figure 2A). The resulting expanded clones demonstrated highly heterogenous me*BRCA1* patterns (Supplementary Figure S5), with one fully methylated and one fully unmethylated epiallele, present at high frequencies and in close to equal proportions (similar to heterozygous methylation of two gene copies, with one methylated and one unmethylated). We chose one heterogeneous heterozygous methylated WEHI-CS62 clone (P4H3), for culture for a further four months and repeated the cell sorting process (Figure 2A). P4H3, as well as the second round FACS clone (P4H3_P3D2) and other second-round clones continued to reveal heterogeneous heterozygous me*BRCA1* patterns (Figure 2B and C; Supplementary Figure S6) after four months in culture (17 passages). Thus, me*BRCA1* patterns were plastic, and continued to generate diversity at the population level, even from single cell clones. However, once fully unmethylated *BRCA1* epialleles were established by DNMT1i treatment, they appeared to be stable for an extended period of time. This raises the concern that loss of silencing of *BRCA1* might be permanent after treatment with DNMT1i.

**Figure 2.**
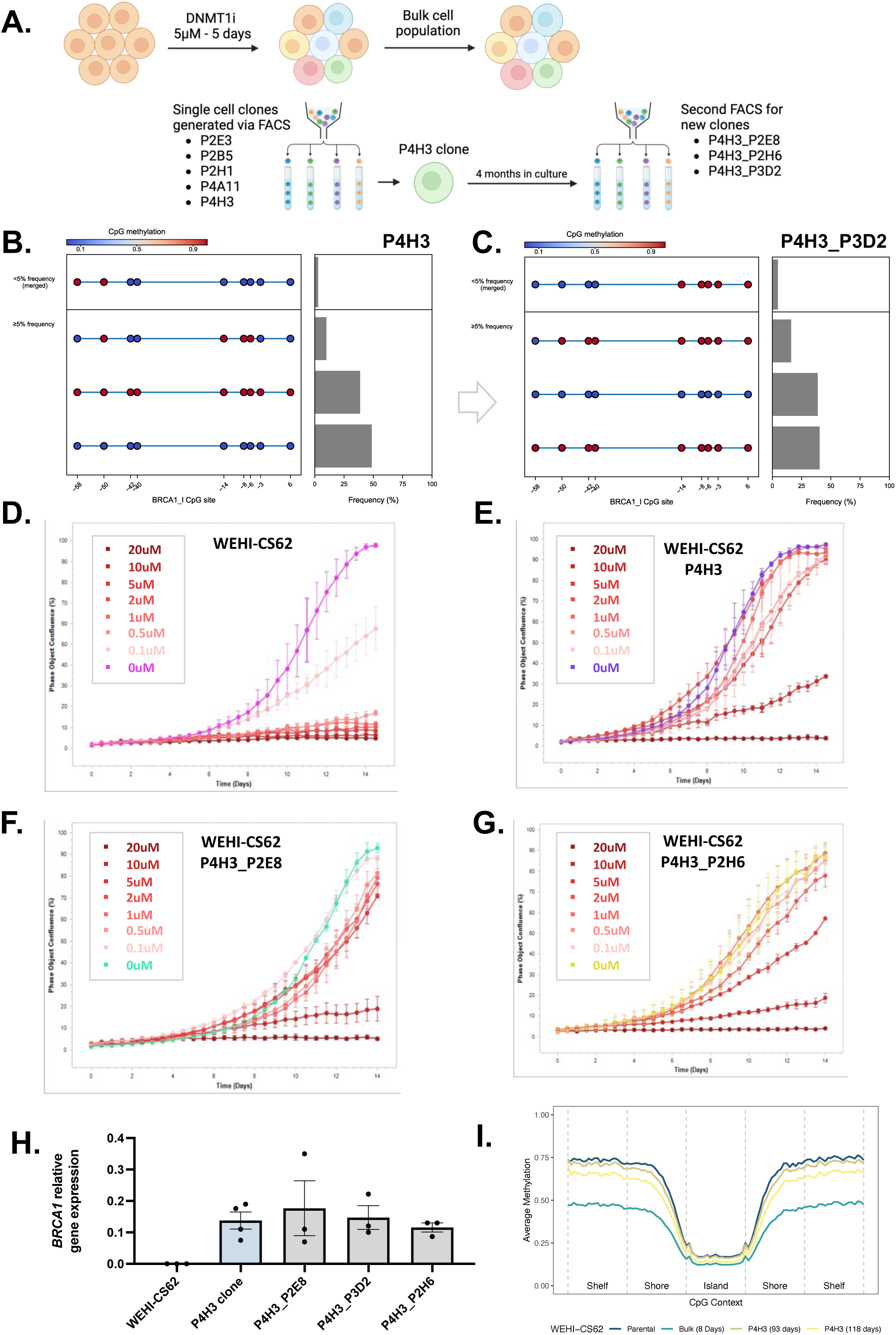
PARPi resistance was observed in WEHI-CS62 clones with me*BRCA1* loss after DNMT1i treatment. (A) Fluorescence-activated cell sorting (FACS) strategy of WEHI-CS62 cells treated with DNMT1i (GSK3484862) for five days. Two days after the end of DNMT1i treatment, single cell clones were generated using FACS. One of these clones (P4H3) with reduced methylation was cultured for another four months (17 passages) before a second FACS sort was performed to generate additional single cell clones, with potentially distinct *BRCA1* promoter methylation patterns. Figure created using BioRender.com. (B) Patterns of me*BRCA1* loss across promoter CpG sites in clone P4H3 were assessed using targeted bisulfite next generation sequencing (meNGS). Individual epialleles present at ≥5% frequency in the sample are presented, along with a merged summary of less frequent epialleles (<5%). Each circle represents CpG sites along the *BRCA1* promoter, and numbers on x-axis indicate CpG distance from *BRCA1* transcription start site. P4H3 demonstrated heterogeneous me*BRCA1*, though fully methylated and unmethylated epialleles were predominant. (C) Similar patterns of me*BRCA1* were detected using meNGS of a clone from the second-round FACS sort (P4H3_P3D2). PARPi rucaparib responses of (D) parental WEHI-CS62 (E) clone P4H3, (F) second-sort clone P4H3_P2E8 and (G) second-sort clone P4H3_P2H6 were assessed over 14 days (cells imaged every 6 hours on Incucyte platform) at the indicated PARPi doses. Mean +/-SEM of two technical replicates presented. Experimental replicates (n=3) presented in Supplementary Figure S8. (H) PARPi-resistant clones (from second-round FACS sort) with reduced me*BRCA1* were found to have increased *BRCA1* gene expression using reverse transcriptase qPCR data. Mean +/-SEM of n=3-4 samples presented, normalized to values for four housekeeping genes. (I) Illumina EPIC global methylation arrays demonstrated global methylation returning to pre-treatment WEHI-CS62 levels across all CpG genomic contexts (averaged CpG values) over time in two post-DNMT1i P4H3 (passage 12) samples.

### DNMT1i treatment drove PARPi resistance in WEHI-CS62 clones

PARPi rucaparib responses were assessed in a series of DNMT1i-treated WEHI-CS62 clones from the first round of cell sorting (described above). All the clones with reduced me*BRCA1* were less responsive to PARPi to various degrees (Supplementary Figure S7). Second generation clones derived from P4H3 were all similarly found to be resistant to PARPi (Figure 2D-G; Supplementary Figure S8-9), with *BRCA1* methylation loss confirmed by meNGS (Supplementary Figure S9-10) and increased *BRCA1* expression observed relative to parental WEHI-CS62 which contained homozygous me*BRCA1* (Figure 2H). Interestingly, global methylation levels in two aliquots of P4H3 passaged for 93-118 days (both 12 passages), appeared to have largely returned to pre-DNMT1i levels of the parental line (Figure 2I) and despite this, *BRCA1* remained unmethylated.

### Global methylation trends in post-DNMT1i WEHI-CS62 samples

Using the Illumina EPIC methylation array (genome-wide methylation analysis tool targeting ∼930K CpG sites) data available for WEHI-CS62 at various timepoints post-DNMT1i, we performed an in-depth analysis of global methylation trends. A differential methylation analysis of probes was performed between two replicates of parental WEHI-CS62 and two replicates of post-DNMT1i clone-P4H3, cultured for 12 passages each (Supplementary Figure S11). Of the 8.5% of probes that were differentially methylated (DM), hierarchical clustering revealed five clusters. While most DM probes lost methylation and did not regain it by 118 days (99% - including *BRCA1*, clusters 1-3), some probes revealed recovering methylation over time (cluster 3), and ∼1% paradoxically gained methylation post-DNMT1i, to levels higher than that observed in the parental cell line (clusters 4-5, with the *MEIS1* gene shown as an example; Supplementary Figure S11-S13). This gain in methylation was mainly observed within the body of genes, consistent with transcriptionally-associated methylation of genes derepressed by loss of promoter methylation (data not shown) [29]. The *BRCA1* promoter did not show restored methylation after four months of P4H3 cell culture (Supplementary Figure S12), supporting our findings above using targeted bisulfite sequencing (Figure 2B and C; Supplementary Figure S6).

### PARPi resistance following DNMT1i treatment was specific to silenced *BRCA1* context

To determine if PARPi resistance caused by DNMT1i treatment was specific to the silenced *BRCA1* context, we utilized the *BRCA2* mutant (PEO1) cell line and a series of OVCAR8 cell line derivatives with various HR defects. In the parental human HGSOC cell line, OVCAR8, heterozygous me*BRCA1* and *BRCA1* gene expression were observed (as described in Supplementary Figure S4), as was PARPi resistance [15]. From two distinct OVCAR8 clones containing landing pad constructs [30, 31], we created one homozygous me*BRCA1* cell line (A6-meth), as well as a *BRCA1* KO cell line (H4-KO), using CRISPR editing, respectively [22]. We also included an OVCAR8 variant with *RAD51C* KO which we have previously described [21]. We focused on these HGSOC lines, as the me*BRCA1* breast cancer cell lines we screened were found to have heterozygous me*BRCA1* (Supplementary Figure S14).

We found that DNMT1i treatment (5 μM GSK3484862 for five days) was associated with PARPi resistance (for both rucaparib and the PARP1-specific PARPi saruparib/ AZD5305), in the homozygous me*BRCA1* cell line, OVCAR8 A6-meth (Figure 3A-D; Supplementary Figure S14-S15). This effect of DNMT1i treatment was not observed in cell lines lacking homozygous me*BRCA1*: the OVCAR8 H4-KO (*BRCA1* KO) cell line; the *BRCA2*-mutant PEO1 cell line; or the OVCAR8 *RAD51C* KO cell line (Figure 3E-F; Supplementary Figure S15-S16). Bisulfite sequencing confirmed loss of me*BRCA1* after DNMT1i treatment in methylated cell lines (Figure 3G; Supplementary Figure S17), and *BRCA1* re-expression in the OVCAR8 A6-meth cell line (Figure 3H). Methylation EPIC array analysis confirmed that DNMT1i treatment had caused global demethylation in all cell lines (Figure 3I). Thus, we observed that treatment with DNMT1i (GSK3484862) caused PARPi resistance only in the context of homozygous me*BRCA1*.

**Figure 3.**
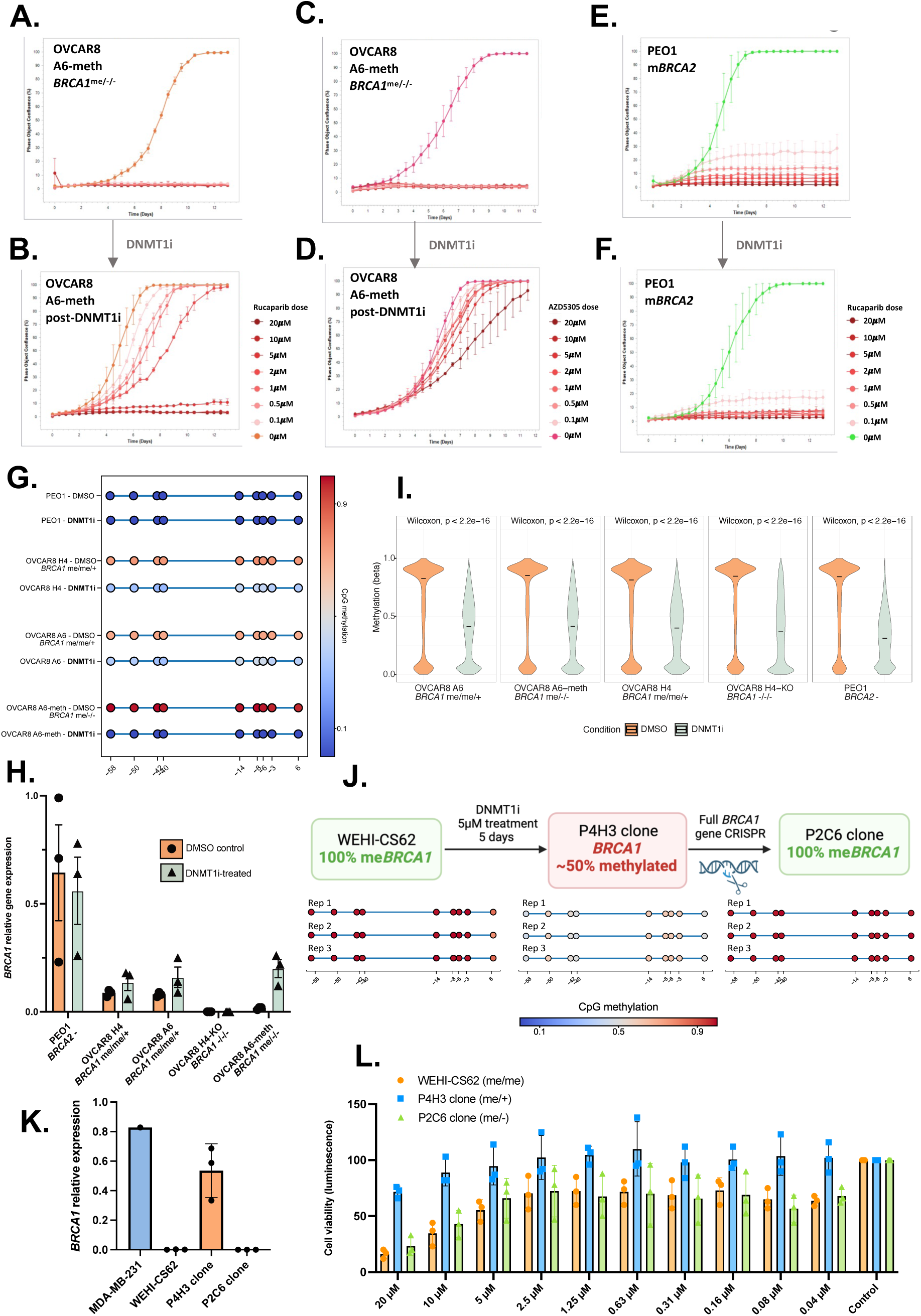
PARPi resistance following DNMT1i was due to loss of *BRCA1* methylation gene silencing. The engineered homozygous me*BRCA1* variant of OVCAR8 (A6-meth, *BRCA1*^me/-/-^) was treated with (A) DMSO control or (B) 5 μM DNMT1i for 5 days. Resulting cells were then treated with indicated doses of PARPi rucaparib and responses were monitored over 14 days using the Incucyte live-cell imaging platform. DNMT1i-treated cells became resistant to rucaparib compared to control (DMSO) cells. The same DMSO (C) or DNMT1i (D) exposed cells were also treated with indicated doses of the potent PARP1-specific PARPi, saruparib (AZD5305), and the same trends were observed over 12 days of treatment. The *BRCA2*-mutant (m*BRCA2*) PEO1 cell line was also exposed to (E) DMSO control or (F) 5 μM DNMT1i for 5 days, and PARPi rucaparib responses to indicated doses assessed over 14 days of treatment. No changes in PARPi response were observed in this cell line. (G) *BRCA1* methylation levels were assessed in all cell lines treated with DMSO or DNMT1i using targeted bisulfite NGS. Each circle represents average methylation level for a CpG site along the *BRCA1* promoter, and numbers on x-axis indicate CpG distance from *BRCA1* transcription start site. (H) *BRCA1* gene expression was also assessed in the same post-DMSO or post-DNMT1i cell line samples using RT-qPCR. *BRCA1* expression normalized to four house-keeping genes. (I) Global DNA methylation changes were assessed using Illumina EPIC methylation arrays in the post-DMSO (control) or post-DNMT1i treated cell line samples, confirming that DNMT1i reduced DNA methylation globally. P values presented (two-tailed Wilcoxon signed-rank test). (J) CRISPR Cas9 gene editing was used to delete a copy of the *BRCA1* gene (including promoter) in DNMT1i-treated WEHI-CS62 clone P4H3, which had ∼50% me*BRCA1*. Of the new single cell clones generated, one (P4H3-P2C6) was found to have homozygous me*BRCA1* (100%). Each circle represents CpG sites along the *BRCA1* promoter, and numbers on x-axis indicate CpG distance from *BRCA1* transcription start site, for n=3 passage replicates (Rep 1-3). (K) P2C6 clone was found to have full *BRCA1* gene silencing by qPCR, like the parental line, and (L) was also re-sensitized to PARPi, as measured by Cell Titre Glo 16 days after saruparib (AZD5305) treatment. Mean +/-SEM of n=3 experiments are presented for H, K and L, and of n=2 technical replicates for A-F (three experimental replicates in Supplementary Figures).

### Restoration of homozygous me*BRCA1* in WEHI-CS62 resensitized cells to PARPi

To confirm that me*BRCA1* loss was indeed the cause of PARPi resistance in cells treated with DNMT1i, we used targeted CRISPR gene editing to specifically KO the re-expressed *BRCA1* gene in DNMT1i-treated WEHI-CS62 cells (clone P4H3 from Figure 2 = *BRCA1*^me/+^). We established a P4H3-subclone with homozygous 100% me*BRCA1* (clone P4H3-P2C6 – *BRCA1*^me/-^; Figure 3J, Supplementary Figure S18). The WEHI-CS62 had two *BRCA1* gene copies [15], and the P4H3 clone was heterozygous, with both a re-expressed and a silenced me*BRCA1* (*BRCA1*^me/+^) allele. Our editing removed the un/hypomethylated copy in this clone (*BRCA1*^me/-^), leaving only one methylated copy of *BRCA1*. Impressively, the resulting fully methylated hemizygous clone, P4H3-P2C6, had complete *BRCA1* gene silencing (Figure 3K) and re-sensitization to PARPi (saruparib/AZD5305), with responses similar to the parental WEHI-CS62 (Figure 3L). Thus, we demonstrated that the impacts of DNMT1i treatment on PARPi responses observed in WEHI-CS62, were dependent on the re-expression of an intact *BRCA1* allele.

### *In vivo* DNMT1i treatment caused *BRCA1* methylation loss

*In vivo* validation experiments were carried out in *BRCA1*-silenced (me*BRCA1*) HGSOC PDX #62, from which the WEHI-CS62 cell line was derived [15]. Tumour fragments from three DNMT1i-treated (45 mg/kg dose, Monday/Wednesday/Friday for four weeks) PDX #62-bearing mice (Figure 4A) were transplanted into six subsequent mice each [15, 23], resulting in 18 PDX-bearing mice (PDX #62-DNMT1i lineage). These were subsequently treated with either rucaparib (450 mg/kg) once, or re-treated with further rucaparib (450 mg/kg) upon tumour progression (described previously [23]), with both vehicle (DPBS control) and untreated controls (Figure 4C). While there was a difference in median time to harvest (TTH) for original PDX #62 treated with rucaparib 450 mg/kg (71 days), compared with the PDX #62-DNMT1i lineage mice treated with the same rucaparib dose (57 days), this difference was not statistically significant (P= 0.811; Supplementary Figure S17; Supplementary Table S4). Fragments from the first three PDX #62 mice treated with DNMT1i (Figure 4A) were harvested upon progression and tested for me*BRCA1* loss using bisulfite sequencing, and no loss was detected (Figure 4C-D; Supplementary Table S5). PDX #62 DNMT1i-treated lineage tumour fragments harvested subsequently, with or without rucaparib treatment, were also tested for me*BRCA1* loss. In one of the three PDX #62 DNMT1i-treated lineages, we observed loss of me*BRCA1*. In this lineage, in one of three mice treated with the PARPi, rucaparib, we observed partial loss of me*BRCA1*. The most significant loss (16% unmethylated epialleles) was observed in a mouse bearing PDX #62 DNMT1i-treated tumour, with no subsequent treatment (Figure 4D-E; Supplementary Table S5). Strikingly, me*BRCA1* loss was associated with restored *BRCA1* gene expression in both mice in which we observed a reduction in me*BRCA1* (Figure 4F).

**Figure 4.**
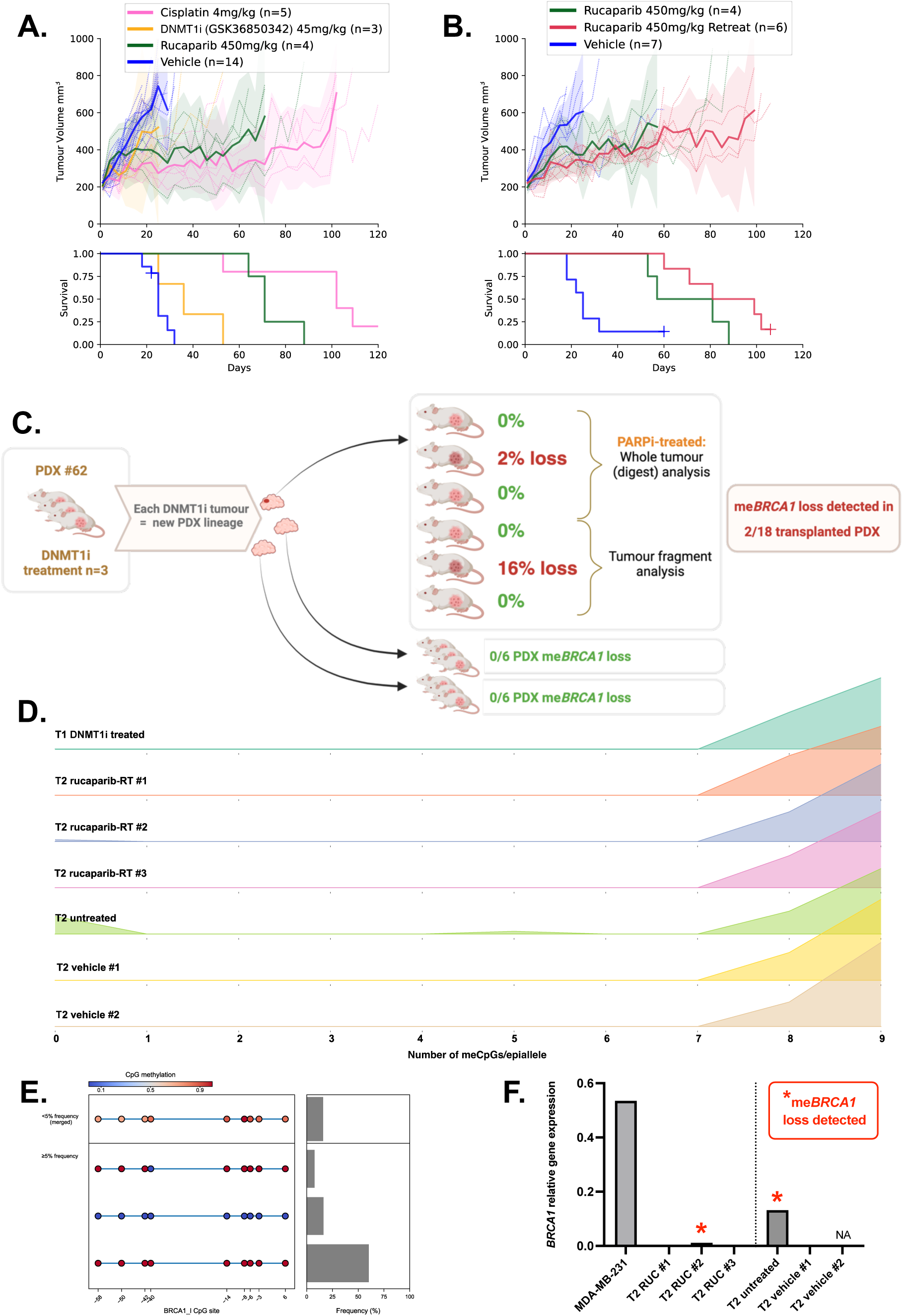
Loss of me*BRCA1* and *BRCA1* re-expression in PDX #62 was observed following *in vivo* DNMT1i treatment. (A) PDX model #62 with stable me*BRCA1* was treated with 45mg/kg of DNMT1i (GSK3685032), 4 mg/kg cisplatin, 450 mg/kg rucaparib or vehicle (DPBS control). DNMT1i treatment produced only a modest therapeutic response compared to PARPi, rucaparib therapy (P= 0.032 and 0.001, for rucaparib and vehicle, respectively). (B) The PDX #62 DNMT1i-treated tumours from (A) were divided and transplanted into new mice to create a new PDX #62-DNMT1i. This lineage was subsequently treated with 450 mg/kg rucaparib single agent, or with 450 mg/kg rucaparib with re-treatment (additional two weeks treatment when tumours reach 500mm^3^) or with vehicle (DPBS control). Time to harvest for the PDX #62-DNMT1i lineage treated with rucaparib 450 mg/kg treatment was numerically reduced compared with the original lineage (57 vs 71 days). Mean PDX tumor volume (mm^3^) ± 95% CI (hashed lines are individual mice) and corresponding Kaplan–Meier survival analysis are shown. Censored events represented by crosses on Kaplan–Meier plot; n = individual mice. Details of time to harvest and Log-Rank test *P* values in Supplementary Table S4. (C) PDX #62 tumours treated with DNMT1i were tested for me*BRCA1* loss using targeted bisulfite sequencing of tumour fragments. No me*BRCA1* loss was detected in the first round of DNMT1i-treated PDX fragments. Second-round PDX #62-DNMT1i lineage transplantation revealed partial loss of me*BRCA1* in two out of six recipients, from one out of three subsequent PDX lineages (0 out of 6 for each of the other two sub-lineages). Figure created using BioRender.com. (D) *BRCA1* meNGS results for the post-DNMT1i lineage where 2/6 mice had detectable me*BRCA1* loss. Y-axis = frequency of each me*BRCA1* epiallele, numbers on x-axis indicate the number of methylated CpG sites per epiallele (0 = fully unmethylated and 9 = fully methylated). T1 = transplant 1, T2 = transplant 2. (E) Detailed *BRCA1* meNGS plot for the PDX tumour with 16% loss of me*BRCA1*, showing the presence of fully unmethylated epialleles. Individual epialleles present at ≥5% frequency in the sample are presented, along with a merged summary of less frequent epialleles (<5%). Each circle represents CpG sites along the *BRCA1* promoter, and numbers on x-axis indicate CpG distance from *BRCA1* transcription start site. (F) *BRCA1* mRNA analysis using RT-qPCR revealed re-expression of *BRCA1* in both PDX tumours with reduced me*BRCA1*: one from a mouse which had not received further treatment (16% me*BRCA1* loss) and one from a rucaparib re-treated (RUC) mouse (2% me*BRCA1* loss).

## Discussion

We have shown that DNMT1 inhibition can drive demethylation of *meBRCA1* and that this demethylation is sufficient to both reverse silencing of *BRCA1* and cause PARPi resistance. Re-expression of a single *meBRCA1* allele is sufficient to restore HR and cause PARPi resistance, as previously described [15]. This demonstrates that DNMT1i therapy may pose a significant risk to patients with epigenetically silenced *BRCA1*, as such treatment can impair an important therapeutically-targetable biomarker and the underlying major mechanism of PARPi response – homozygous me*BRCA1*. PARPi therapy has transformed the treatment outlook for some women with *BRCA*-mutated/HRD HGSOC, but the importance of our observations for me*BRCA1*/HRD HGSOC needs to be explored in existing clinical (trial) samples, as post-PARPi resistance in the clinic is often profound, can involve cross-resistance with platinum and is, as yet, poorly understood.

The observed ability to reverse gene silencing with DNMT1i suggests that *BRCA1* silencing in cancer does not involve robust multilayered changes in chromatin state or active reinforcement and maintenance, relying only on propagation of the methylation state at replication by DNMT1. Our experiments also demonstrate ongoing selection for *BRCA1* expression, even without subsequent therapy, in both our cell culture and PDX models. Notably, *BRCA1* expression was maintained following DNMT1i treatment, despite most other CpG sites returning to pre-DNMT1i levels of methylation, following four months of culture. The extreme rarity of stably homozygous me*BRCA1* and *BRCA1*-silenced cell line models, further supports the concept of cell culture selection for *BRCA1* expression. Given that DNMT1i could reverse silencing in the uniquely me*BRCA1*-stable WEHI-CS62 cell line, we hypothesise that this ability to reverse gene silencing could apply generally to silenced *BRCA1* in cancer.

Furthermore, we believe our findings are likely to be relevant to other genes of the HR pathway, and other cancer types. Homozygous promoter methylation of the *RAD51C* gene (me*RAD51C*) is also a known PARPi-response biomarker in HGSOC (found in ∼3.7% of cases) [32], with loss of methylation associated with PARPi resistance [23]. Unfortunately, a lack of HGSOC cell lines with me*RAD51C* has restricted our ability to study the effects of DNMT1i in this context. Similarly, a lack of breast cancer cell lines with homozygous me*BRCA1* has also limited our ability to validate findings in triple negative breast cancer (TBNC), where me*BRCA1* is frequently observed (∼15% of cases), and where loss of me*BRCA1* is also associated with restored HR repair [33]. Given that PARPi therapy is already approved for breast cancers with inherited *BRCA1/2* pathogenic variants, and DNMTi have been investigated in breast cancer trials (NCT04134884; NCT00748553; NCT01349959), our findings have important potential relevance for this common cancer type. There may even be risk for other cancer types where me*RAD51C* is observed, such as in stomach adenocarcinoma (∼6% of cases) [32], where previous work has demonstrated increased *RAD51C* expression following *in vitro* treatment of me*RAD51C* cells with decitabine [34].

While inhibiting the maintenance of methylation is sufficient to reactivate silenced *BRCA1,* the generation of single cell clones that have ongoing instability post treatment suggests that this is not a single step process. While the methylation state within the sequenced region was mainly homogenous, propagation of methylation across the broader promoter region may account for this variability across multiple replication cycles. Additionally, the methylation at the silenced *BRCA1* promoter may be directing changes in chromatin state, both directly and indirectly through transcriptional inactivity. As silencing is removed, we hypothesise this transcriptional activity is self-reinforcing in the absence of an active silencing mechanism, promoting an active chromatin state. Histone deacetylase inhibitors (HDACi) are known to enhance the demethylating effects of DNMTi [35, 36], but have also been shown to cause DNA demethylation alone [35, 37–39]. These findings support a role for transcriptionally active acetylated chromatin in the erasure of DNMT1 maintained methylation marks, and suggests similar destabilisation of silencing and re-expression of *BRCA1* may be possible with HDACi.

Loss of me*BRCA1* was detected in two of six PDX #62 tumours in a lineage transplanted following *in vivo* DNMT1i treatment. Rates of me*BRCA1* loss may have even been higher, given our methylation measurements of most (12/18) tumours were via tumour-fragment based analysis, where tumour heterogeneity would impact detection. Indeed, me*BRCA1* loss was not detected in the tested fragment of the DNMT1i-treated tumour which yielded two subsequent tumours where me*BRCA1* loss was detected. Given our ability to detect loss of silencing in a small total tumour volume after short duration treatment at clinically relevant doses, our *in vivo* results clearly indicate that loss of silencing is likely to occur in at least some patients treated with DNMT1i. As our work was conducted in NOD-scid IL2rγnull (NSG) mice which lack a T cell response, we could not address the immunogenic impacts of DNMT1i relative to PARPi in the treatment of HGSOC.

In conclusion, we have provided compelling evidence that DNMT1i is able to reverse *BRCA1* methylation and silencing, and that even partial loss of me*BRCA1* in this context results in restored *BRCA1* expression and PARPi resistance. Using PDX, we demonstrated *in vivo* that DNMT1i could cause loss of me*BRCA1*. Thus, DNMT1i pose a risk for patients with HGSOC with homozygous *BRCA1* methylation, the second largest group of HGSOC patients likely to respond to PARPi (after *BRCA1/2* mutated cases). For cancer patients, DNMT1i places their chance of having a durable response to PARPi, and/or subsequent platinum-based therapy, at risk. These negative impacts of exposure to DNMT1i should be considered for patients with known *BRCA1* methylation in their cancer, and could be explored in existing HGSOC clinical trial samples exposed to DNMT1i. In cancer types with a high frequency of silenced *BRCA1* (such as HGSOC), this caution should extend to non-*BRCA1/2* mutated HRD cases lacking methylation information.

## Acknowledgements

We thank Silvia Stoev, Kathy Barber, Chloe Neagle, Steph Bound and Dan Fayle for technical assistance. This work was made possible through the Australian Cancer Research Foundation, the Victorian State Government Operational Infrastructure Support and Australian Government NHMRC IRIISS. AOCS would like to thank all of the women who participated in these research programs. The AOCS would also like to acknowledge the contribution of the study nurses, research assistants, and all clinical and scientific collaborators to the study. The complete AOCS Study Group can be found at www.aocstudy.org. The Australian Ovarian Cancer Study gratefully acknowledges additional support from Ovarian Cancer Australia and the Peter MacCallum Foundation.

## Consortia

Australian Ovarian Cancer Study (AOCS), G. Chenevix-Trench, A. Green, P. Webb, D. Gertig, S. Fereday, S. Moore, J. Hung, K. Harrap, T. Sadkowsky, N. Pandeya, M. Malt, A. Mellon, R. Robertson, T. Vanden Bergh, M. Jones, P. Mackenzie, J. Maidens, K. Nattress, Y. E. Chiew, A. Stenlake, H. Sullivan, B. Alexander, P. Ashover, S. Brown, T. Corrish, L. Green, L. Jackman, K. Ferguson, K. Martin, A. Martyn, B. Ranieri, J. White, V. Jayde, P. Mamers, L. Bowes, L. Galletta, D. Giles, J. Hendley, T. Schmidt, H. Shirley, C. Ball, C. Young, S. Viduka, H. Tran, S. Bilic, L. Glavinas, J. Brooks, R. Stuart-Harris, F. Kirsten, J. Rutovitz, P. Clingan, A. Glasgow, A. Proietto, S. Braye, G. Otton, J. Shannon, T. Bonaventura, J. Stewart, S. Begbie, M. Friedlander, D. Bell, S. Baron-Hay, A. Ferrier (decd.), G. Gard, D. Nevell, N. Pavlakis, S. Valmadre, B. Young, C. Camaris, R. Crouch, L. Edwards, N. Hacker, D. Marsden, G. Robertson, P. Beale, J. Beith, J. Carter, C. Dalrymple, R. Houghton, P. Russell (decd.), M. Links, J. Grygiel, J. Hill, A. Brand, K. Byth, R. Jaworski, P. Harnett, R. Sharma, G. Wain, B. Ward, D. Papadimos, A. Crandon, M. Cummings, K. Horwood, A. Obermair, L. Perrin, D. Wyld, J. Nicklin, M. Davy, M. K. Oehler, C. Hall, T. Dodd, T. Healy, K. Pittman, D. Henderson, J. Miller, J. Pierdes, P. Blomfield, D. Challis, R. McIntosh, A. Parker, B. Brown, R. Rome, D. Allen, P. Grant, S. Hyde, R. Laurie, M. Robbie, D. Healy, T. Jobling, T. Manolitsas, J. McNealage, P. Rogers, B. Susil, E. Sumithran, I. Simpson, K. Phillips, D. Rischin, S. Fox, D. Johnson, S. Lade, M. Loughrey, N. O’Callaghan, W. Murray, P. Waring, V. Billson, J. Pyman, D. Neesham, M. Quinn, C. Underhill, R. Bell, L. F. Ng, R. Blum, V. Ganju, I. Hammond, Y. Leung, A. McCartney (decd), M. Buck, I. Haviv, D. Purdie, D. Whiteman & N. Zeps

**AOCS consortium representative contact:**

