## Supplementary Methods for "Demethylating agents drive PARP inhibitor resistance in ovarian carcinomas with *BRCA1* gene silencing"

### De-methylating agents drive methylation loss and PARP inhibitor resistance in ovarian carcinomas with *BRCA1* gene silencing

Nesic et al., 2024

**EPIC methylation arrays and analysis**

Genome-wide DNA methylation arrays were performed on bisulfite converted DNA using the Infinium MethylationEPIC v2.0 Kit (Illumina) at the Australian Genome Research Facility (AGRF).

The raw intensity files were converted into beta values using minfi (version 1.44.0), followed by background correction and normalisation with function preprocessIllumina. Probes that are either non-autosomal, dropped out in measurement, had no intensity, had beads count less than 3 or had a detection p value less than 0.01 were removed from down-stream analysis. The differentially methylated regions were identified for WEHI-CS62 post-GSK treatment (pre-GSK treatment N =2, long-term recovered N =3) and CpGs were annotated using information from NCBI and EBI (MANE) (GRCh38, v1.3).The raw intensity files were converted into beta values using minfi [1] (version 1.44.0), followed by background correction and normalisation with function preprocess Illumina. Probes that are either non-autosomal, dropped out in measurement, had no intensity, had bead counts less than 3 or had a detection p-value less than 0.01 were removed from downstream analysis. Further filtration removed flagged probes using the Infinium MethylationEPIC v2.0 data (EPIC-8v2-0_A1-FlaggedProbes.csv, downloaded June 14, 2023). Differentially methylated probes (DMPs) were identified for WEHI-CS62 (pre-DNMT1i treatment, N = 2; long-term recovered, N = 3) from M values (logit base 2 conversion from beta values) using the R package limma [2] (version 3.54.2). Batch effects were accounted for in the linear regression model (one pre-treated and one recovered sample were assessed in one batch, the rest in a separate batch). DMPs were filtered based on a beta value change of ≥ 0.25.

Hierarchical clustering of DMPs was performed using R package fastcluster (version 1.2.3) and plotted using ComplexHeatmap [3] (version 2.14.0). Significance was determined by Benjamini-Hochberg corrected p-values of less than 0.05. The CpGs were annotated using information from NCBI and EBI (MANE) (GRCh38, v1.3).

**Generation of WEHI-CS62 Cas9 expressing cell line for CRISPR screen**

HEK 293T cells were cultured in FV media (without penicillin streptomycin), before being trypsinized and plated 6 x $10^6$ cells in 12ml per 10cm plate, making 2 plates. These cells were then incubated at 37°C in 5% CO2 overnight, allow cells to settle. The following day, 41µl of Lipofectamine 3000 Transfection Reagent (Thermo Fisher Scientific, Cat# L3000008) was added to 1.5ml of OptiMEM media (Gibco, Cat# 31985070) to create reaction "mix A". Then a separate reaction "mix B" was made by adding 35µl of p3000 (Thermo Fisher Scientific, cat# L3000008) to 1.5ml of OptiMEM media (Gibco, Cat# 31985070) and reagents in Methods Table 1 for packaging FUCas9Cherry (Cas9 and mCherry) plasmid DNA or Methods Table 2 for packaging Brunello plasmid DNA. FUCas9Cherry plasmid DNA (Addgene #70182), viral component plasmids pMDL (Addgene #12251), VSVg (Addgene #12259) and RSV-REV (Addgene #12253) were provided by the Herold Lab (WEHI). Human CRISPR Knockout Pooled Library Brunello plasmid DNA (Addgene Cat# 73178) was provided by E. Surgenor of the Huntington group (WEHI). To create the plasmid DNA: lipid complex, 1.5ml of mix A was mixed with 1.5ml of mix B to create "mix C". This mixture was then incubated for 10-20 minutes at room temperature. HEK 293T cells were then transfected with the packaged lentivirus. After incubation, 6ml of FV media from each HEK 293T cell dish was removed and 3ml of mix C was added to each dish.

Cells were then incubated for 6 hours at 37°C in 5% CO2. The media-virus mixture was then removed and replaced with 12ml of regular FV media and cells incubated at 37°C in 5% CO2 for 24 hours. Packaged lentivirus was harvested at 24 hours post-transfection. All viral supernatant (12ml) was removed and replaced with fresh FV media (12ml). The viral supernatant was centrifuged at 800g for 10 minutes at room temperature to remove cellular debris. The resulting clarified supernatant was then filtered through a 45µm pore to remove any remaining cellular debris. This material was stored at 4°C. At 52 hours post-transfection, this process was repeated, for a total of 24ml of packaged virus produced per dish of infected cells. The virus was dispensed as 5ml aliquots in 10ml tubes and stored in the -80 freezer until required for infections.

**Methods Table 1. Reagents used for lentiviral packaging of FUCas9Cherry plasmid DNA.**

| Reagent | Concentration (µg/µl) | µg required per mix B reaction | µl added per mix B reaction |
|---|---|---|---|
| pMDL | 1.4 | 5 | 3.6 |
| RSV-REV | 0.85 | 2.5 | 2.9 |
| VSVg | 2.3 | 3 | 1.3 |
| Plasmid DNA (Cas9) | 1.7 | 5 | 2.9 |

**Methods Table 2. Reagents used for lentiviral packaging of Brunello plasmid DNA.**

| Reagent | Concentration (µg/µl) | µg required per mix B reaction | µl added per mix B reaction |
|---|---|---|---|
| pMDL | 1.4 | 5 | 3.6 |
| RSV-REV | 0.85 | 2.5 | 2.9 |
| VSVg | 2.3 | 3 | 1.3 |
| Plasmid DNA (Brunello) | 2 | 5 | 2.5 |

For infections, previously prepared Cas9-packaged virus was thawed, and 2 x $10^5$ cells were plated in each well of 24-well plate in 500µl (4 replicate wells). Then 8µg/mL polybrene was added to neat viral solution, and 1ml of this solution was added to each well with cells. An uninfected control plate was also prepared, where FV media was added instead of virus. All plates were then centrifuged at 800g for 1 hour at 32°C to spin infect cells. Plates were then incubated overnight at 37°C in 5% CO2. The following day, virus/media was removed and cells were washed twice in DPBS. Fresh FV media was then added to each well and plates were incubated overnight at 37°C in 5% CO2. Cells were then washed again in DPBS, detached using trypsin, washed again using DPBS, and sorted for mCherry fluorescence on the BD FACSAria™ III sorter; uninfected/negative control cells (with and without DAPI stain), and infected mCherry positive cells collected in 1ml FCS. Resulting mCherry/Cas9 positive cells were then cultured for 6-7 days before repeating the Fluorescence Activated Cell Sorting (FACS) protocol (second sort to enrich further for Cas9-positive cells).

**WEHI-CS62 Brunello CRISPR screens**

In order to find the optimal concentration of puromycin to use in selection of Cas9-expressing cells with guides in the Brunello CRISPR screen (i.e. a concentration that is not overly toxic and just enough to kill all uninfected cells to avoid creating bias in the screen), toxicity curves were generated for each cell line. Uninfected cas9 cell lines were plated onto two (duplicate) 12-well cell culture plates using their standard split-rate (adjusted for change in surface area) for a total volume of 2ml. Cells were then incubated at 37°C in 5% CO2 overnight, allow cells to settle. The following day, puromycin (Invivogen, Cat# ant-pr-1), diluted to 100µg/ml in DPBS, was added to the 2ml of FV media per well to create final puromycin concentrations of 1, 0.9, 0.8, 0.7, 0.6, 0.5, 0.4, 0.3, 0.2, 0.1 and 0µg/ml. Wells were checked under 4x and 10x magnification after 3 days to assess cell death in each well. The lowest puromycin concentration causing complete cell death in the well was assessed visually, and used as the selecting concentration in the Brunello CRISPR screen for each cell line.

Infection efficiency represents the percentage of infected cells per reaction. To uncover the dilution of Brunello virus required for a 30-50% infection efficiency for the CRISPR PARPi resistance screen, 2 x $10^5$ cells were plated in each well of a 24-well plate in a volume of <100µl. A lentiviral dilution series containing 8µg/mL polybrene was created: neat, 1:1, 1:2, 1:3, 1:4, 1:5, 1:10, 1:15, 1:20, 1:30, 1:40, no virus. 1.5 mL of each lentiviral dilution was added per well with cells. Cells were spin infected by centrifugation at 800g for 1 hour at 32°C. Plates were then incubated overnight at 37°C in 5% CO2. The following day, the virus was removed and cells were washed twice with DPBS. Fresh FV media was added to the cells and plates were incubated overnight at 37°C in 5% CO2. The following day, cells were washed with DPBS again, then trypsinized and counted. 2000 cells were seeded in 75µl per well on a CELLSTAR® 96 well plate (Greiner, Cat# M1062), with 4 wells seeded per lentiviral dilution/condition. Cells were left to settle at 37°C in 5% CO2 for 1 hour before adding 75µl at 1.4µg/ml of puromycin in FV media to 2 wells, and adding 75µl of regular FV media to the other 2 wells (control). Plates were incubated at 37°C in 5% CO2 for 3 days. CTG assay was then used to measure the viability of the cells, and MOI calculated by dividing the infected cell viability by the control cell viability for each lentiviral dilution.

**Brunello library CRISPR infections**

Cas9-expressing versions of each cell line were expanded in Millicell HY 5-layer cell culture T-1000 flasks (MERCK, Cat# PFHYS1008) to produce enough cells for Brunello CRISPR guide library infections. Once expanded, 70 million cells were dissociated using trypsin and resuspended in 96mL of FV media with 1:40 diluted Brunello virus containing 8µg/mL polybrene (2.4ml thawed virus, 76.8µl of 10mg/ml polybrene). These cells were plated across 16 x 6-well plates (2ml per well) per infection replicate/experiment. Also, 4.3 million infected/uninfected cells were plated in a T75 flask each as puromycin selection controls. All infection plates were then centrifuged at 800g for 1 hour at 32°C to spin infect cells (8 plates per spin, two spins required). Plates were then incubated overnight at 37°C in 5% CO2. The following day, virus/media was removed and cells were washed twice in DPBS before being dissociated and total cells transferred into multilayer flasks for cell culture. Puromycin selection was then performed (24 hours post infection) using 1.4µg/mL. At least 30 million cells were maintained per passage for 7 days of puromycin selection/culture.

Total cells were then harvested and 2 cell pellets of 20 million cells each were snap frozen for DNA extraction and library diversity assessment. Remaining cells were used to make viable DMSO (10%) freezings of 25 million cells per vial, and these would be thawed for downstream CRISPR screen rucaparib treatment experiments.

**Brunello CRISPR screen drug treatments**

Cas9-expressing WEHI-CS62 cells were infected with the Brunello library, which comprises of 77,441 single guide RNAs (sgRNAs; an average of 4 sgRNAs per gene), and 1000 non-targeting control sgRNAs. For the WEHI-CS62-Cas9 cell line, 70 million cells were infected using a 50% infection efficiency, to result in >30 million cells with an sgRNA following puromycin selection to remove uninfected cells, providing sufficient diversity for this CRISPR library (~385x sequencing coverage per guide [4]).

25 million WEHI-CS-62 Cas9 Brunello cells from each infection were thawed and expanded, so that 12.5 million cells were plated per treatment/control flask (2 replicates per infection). WEHI-CS-62 Cas9 Brunello cells were then treated with escalating doses of rucaparib (EC50 of 1.2µM for 4 weeks and then EC75 5µM for 4 weeks) or DMSO (matched to rucaparib concentration, harvested at 7 weeks due to higher confluence). Escalating doses were used to prevent excessive cell death, given that these assays were run longer than the CTG experiments

used to calculate the used EC50 and EC75 values. DMSO control flasks (equivalent volume to DMSO added in rucaparib-treated flasks) were run in parallel, to be used in the final analysis.

A minimum of 12.5 million cells were maintained per passage. 15 million cells from each treatment were harvested and snap frozen for DNA extraction (Zymo Quick-DNA Midiprep plus kit; Cat #D4075) and library preparation. DNA was then bisulfite converted and me*BRCA1* analysed by ddPCR (as described in previous sections).

**CRISPR screen library generation and sequencing**

A single-step PCR was used to create Brunello NGS libraries using primers outlined in Table 2.7. DNA was diluted to 100ng/µl and 15µl was added to 50µl of Taq 2X Master Mix (NEB; cat# M0270L), 5µl of each 10µM primer dilution (Table 2.7) and 25µl of molecular grade H2O. Reactions were incubated at 95°C for 1 minute, followed by 25 cycles of 95°C for 30 seconds, 53°C for 30 seconds, 72°C for 30 seconds. Reactions then incubated at 72°C for 10 minutes, and stored at 4°C. Resulting libraries were cleaned up using a 0.9:1 Ampure XP beads to PCR product ratio (45µl beads into 50µl PCR product). Libraries were then quantitated using the Qubit dsDNA HS assay (Invitrogen, Cat#Q32854), and the size of each library was assessed using the Agilent D1000 ScreenTape System according to manufacturer's instructions (ScreenTape: Cat# 5067- 5582, D1000 Reagents: Cat# 5067- 5583). Libraries were pooled equally based on molarity (calculated using DNA concentrations and average library size). Pooled libraries were sequenced on the Illumina Nextseq500 platform by the WEHI genomics hub using a TG NextSeq® 500/550 v2 High Output Kit (75 cycles; Cat# TG-160-2005). Resulting FASTQ files were run through a custom in-house pipeline to report all detected unique Brunello guide sequences, their corresponding genes and read counts.

**CRISPR screen analysis**

To generate a heatmap of *DNMT1* and *DNMT3A* guide counts before and after drug treatment in each sample, the R software package edgeR v3.40.2 was used to extract the counts per million for each guide, an offset of 1 was added, before the data were $\log_2$ transformed and plotted using the R package pheatmap v1.0.12. A bar plot was generated using R v4.2.2 to visualize the number of sgRNAs from the Brunello library detected across all sequenced samples, in each infection before treatment, and after drug or DMSO treatment in each sample in the CRISPR screen.
