## Supplementary Figures for "Demethylating agents drive PARP inhibitor resistance in ovarian carcinomas with *BRCA1* gene silencing"

### De-methylating agents drive methylation loss and PARP inhibitor resistance in ovarian carcinomas with *BRCA1* gene silencing

Nesic et al., 2024

### Figure S1

A.

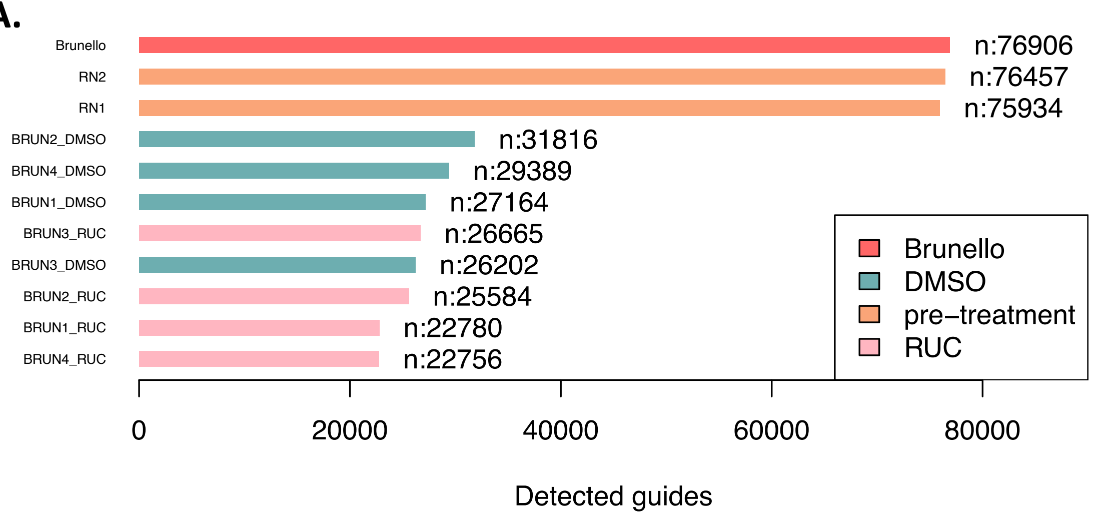


B.

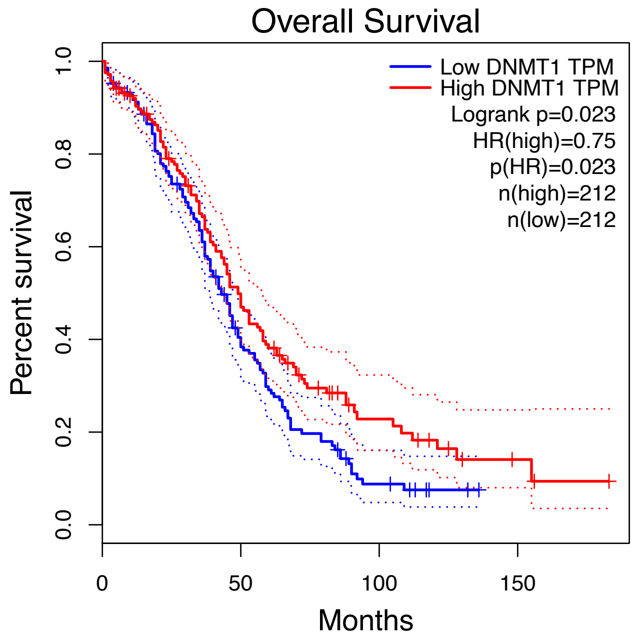


**Supplementary Figure S1. *DNMT1* is an essential gene that might impact HGSOC survival outcomes.**

**A.** Of the 22,756 to 26,665 guides detected post-PARPi at the end of a genome-wide PARPi resistance CRISPR screen of WEHI-CS62, *DNMT1* guides were not detected. “Brunello” shows the total guides contained in the CRISPR screen library, “DMSO” indicates control treatment arms, “RUC” indicates the rucaparib-treated arms and “pre-treatment” indicates post-infection but pre-treatment samples used to test for library diversity, where RN1 is infection replicate 1 and RN2 is infection replicate 2. **B.** The Gene Expression Profiling Interactive Analysis (GEPIA) was used to interrogate the impacts of *DNMT1* gene expression on survival outcomes in ovarian cancer using the TCGA ovarian cancer expression and survival data [1]. A Kaplan-Meier plot of overall survival is presented. A median group cut-off of 50% was used to categorize expression as low or high. 95% confidence interval is indicated by dotted lines. Hazards ratio based on Cox PH Model.

**Figure S2**

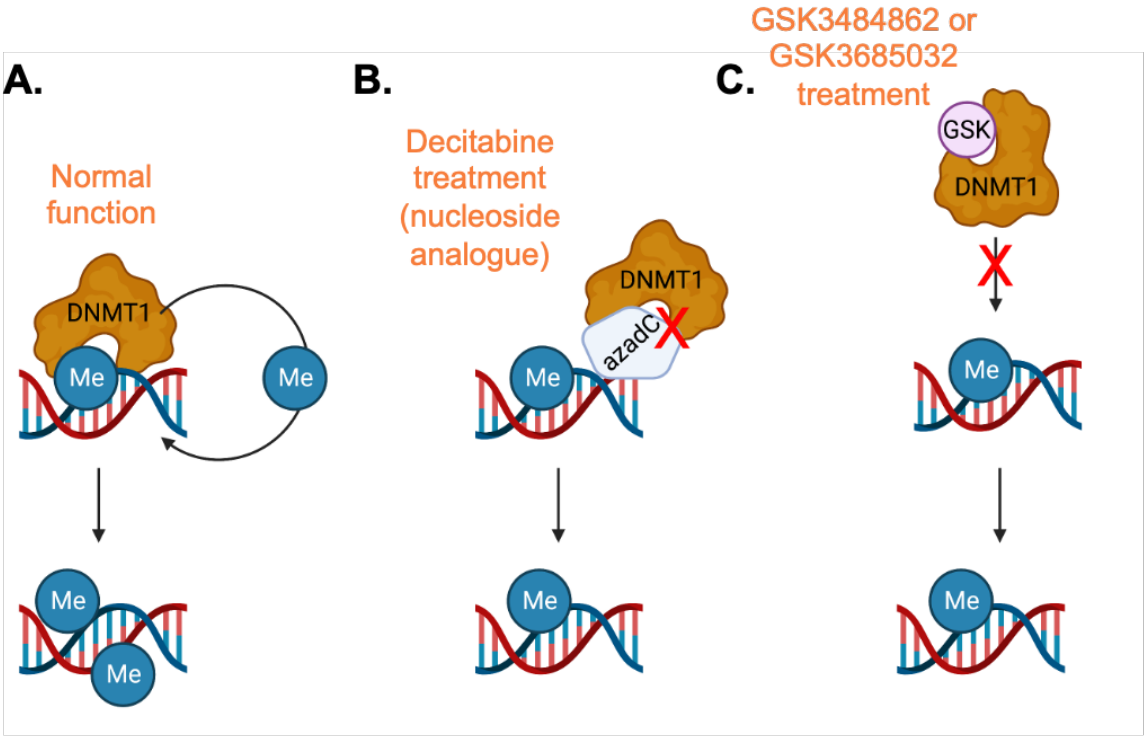


**Supplementary Figure S2. DNMT1-targeting compounds and their mechanisms of action.**

**A.** Under normal conditions, DNMT1 maintains DNA methylation patterns by binding to hemi-methylated DNA and adding a methyl group to the complementary newly synthesized unmethylated DNA strand. **B.** Decitabine is a nucleoside analogue that traps DNA methyl-transferases including DNMT1 on the DNA, thus inhibiting maintenance of DNA methylation following DNA replication, as well as de-novo DNA methylation (by trapping DNMT3A, for example). **C.** Decitabine thus has a broader set of effects compared to DNMT1-specific inhibitors like GSK3484862 and GSK3685032, which inhibit the catalytic site of DNMT1 to prevent its DNA-methylating action. Figure was generated using BioRender.com.

### Figure S3

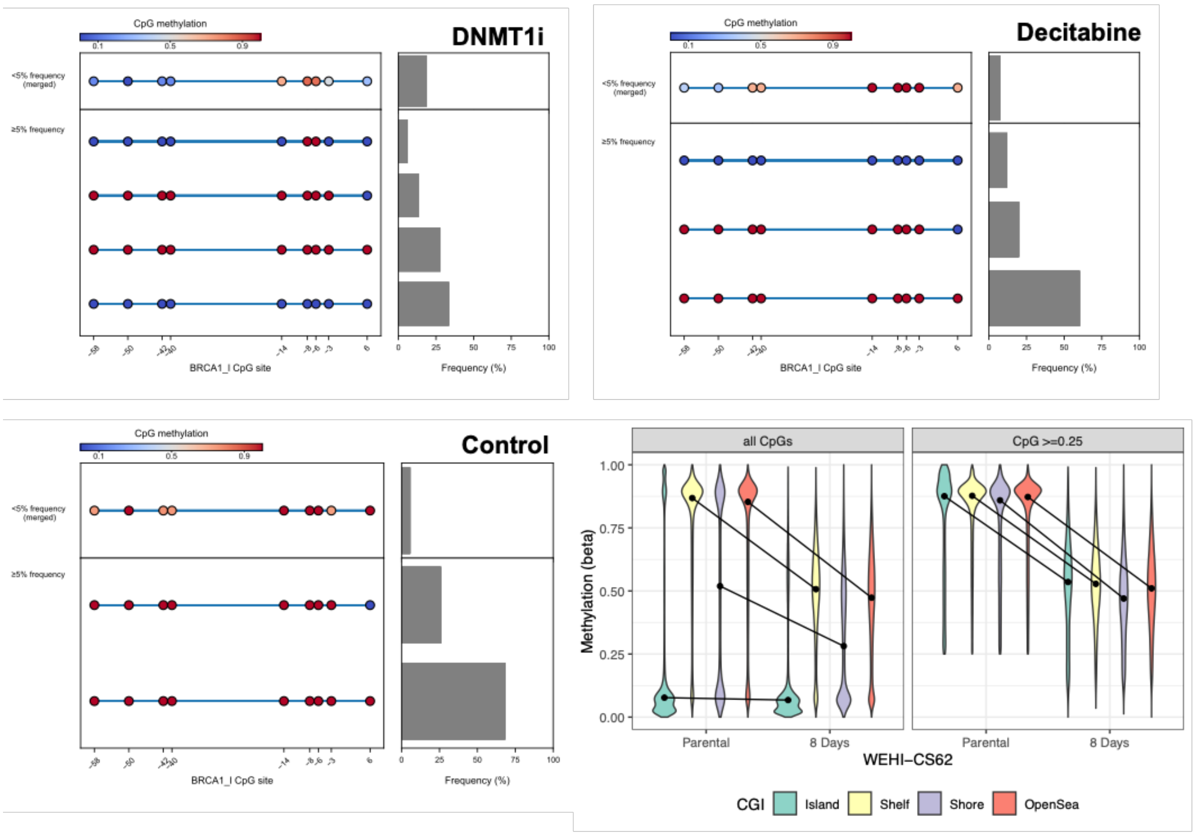


**Supplementary Figure S3. Loss of me*BRCA1* and global methylation observed in WEHI-CS62 post-DNMTi treatment.**

The presence of fully unmethylated epialleles was confirmed in **A.** GSK3484862 10uM, and **B.** decitabine 0.1uM treated WEHI-CS62 samples, while these were not observed in **C.** DMSO control. Patterns of me*BRCA1* loss across promoter CpG sites were assessed using targeted bisulfite next generation sequencing (meNGS) of the *BRCA1* promoter. Individual epialleles present at ≥5% frequency in the sample are presented, along with a merged summary of less frequent epialleles (<5%). Each circle represents CpG sites along the *BRCA1* promoter, and numbers on x-axis indicate CpG distance from *BRCA1* transcription start site. **D.** Illumina EPIC methylation arrays were used to confirm global decreases in DNA methylation in WEHI-CS62 following GSK3484862 10uM treatment. Each violin represents the distribution of global methylation with median value marked in circle. The trend of change induced by DNMT1i in median value were highlighted with line. Changes in CpGs that were methylated in the untreated parental (beta value ≥ 0.25) were not restricted to any particular CpG contexts, such as CpG island (green), shelf (yellow) shore (purple) or open sea (red).

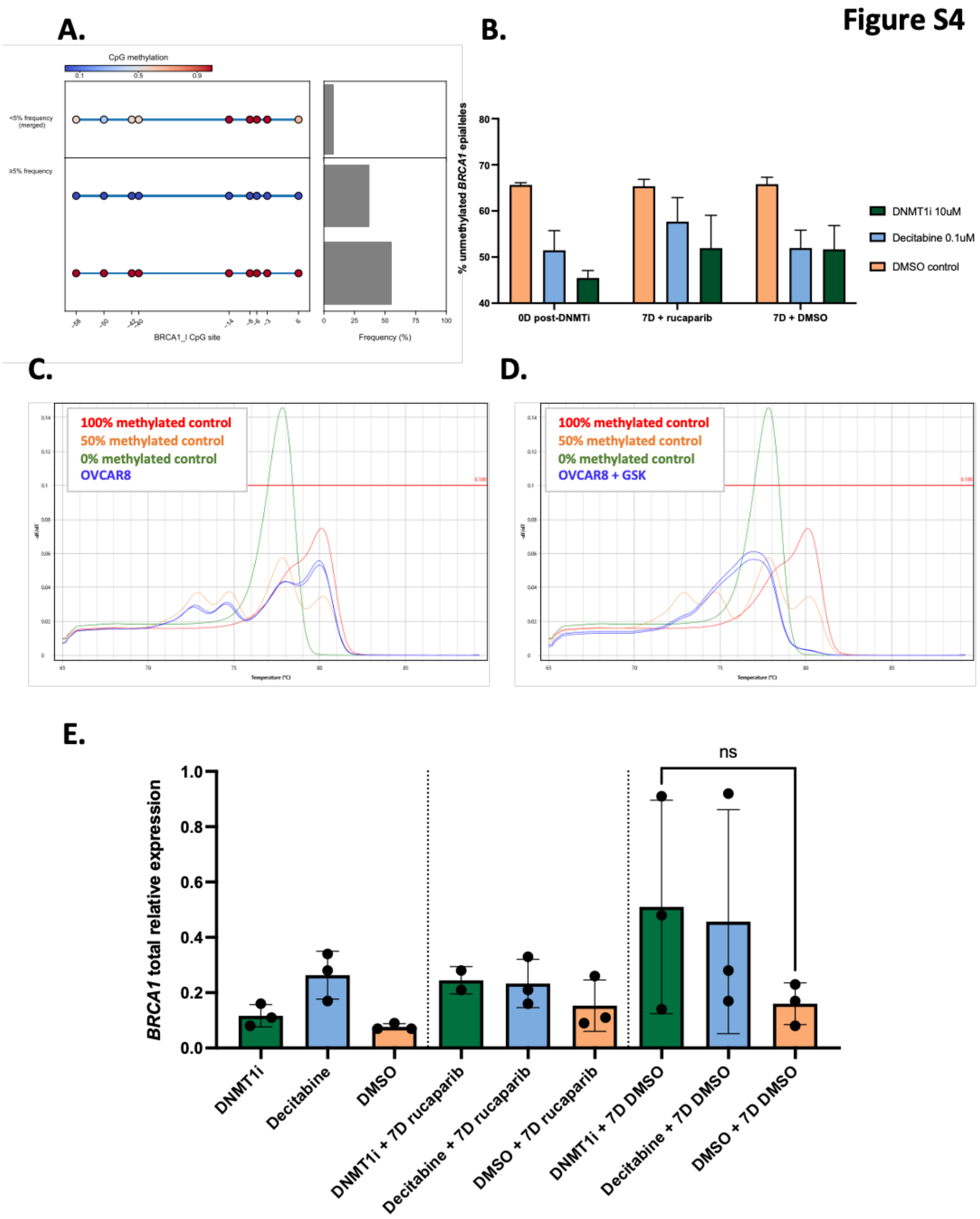


**Supplementary Figure S4. DNMTi caused methylation loss in a cell line with heterozygous me*BRCA1*.**

**A.** OVCAR8 has heterozygous me*BRCA1* patterns (two methylated and one unmethylated copy), assessed using targeted bisulfite next generation sequencing (meNGS) of the *BRCA1* promoter. Individual epialleles present at ≥5% frequency in the sample are presented, along with a merged summary of less frequent epialleles (<5%). Each circle represents CpG sites along the *BRCA1* promoter, and numbers on x-axis indicate CpG distance from *BRCA1* transcription start site. **B.** Loss of me*BRCA1* was observed following DNMT1i (GSK3484862/

GSK) and Decitabine treatment of OVCAR8, measured using ddPCR and **C.** MS-HRM of OVCAR8 cell line. Each line represents a measurements per tumor/sample (indicated in colour legend). RFU, relative fluorescence units. Y-axis is the derivative of fluorescence over temperature [Δd(RFU)/dT] versus temperature (T). The red horizontal line is a default threshold set by the software to report peak melting temperatures, and can be manually changed. It was left at default setting of 0.01, as it is not relevant for this type of analysis. **D.** MS-HRM of OVCAR8 following treatment with GSK3484862 (DNMT1i). **E.** RT-qPCR of *BRCA1* mRNA indicated heterogeneous increases in *BRCA1* expression 7 days (7D) after DNMT1i/decitabine + DMSO treatment in OVCAR8 cells (not significant). Each dot represents value for one of N=3 replicate experiments, however one DNMT1i +7D rucaparib outlier was removed. Mean ± SEM is presented.

Figure S5

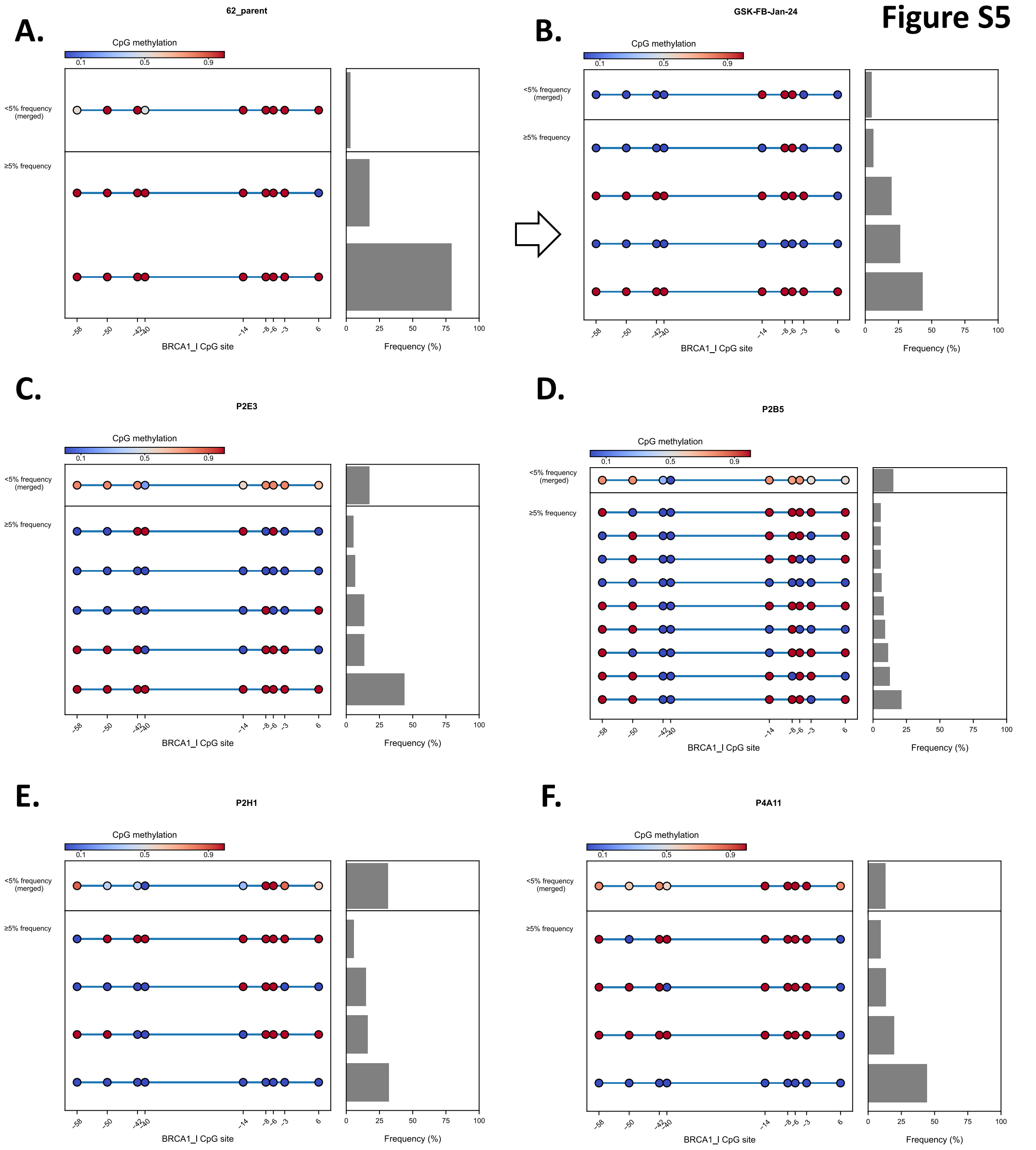


**Supplementary Figure S5. DNMT1i caused heterogenous *BRCA1* methylation loss in individual WEHI-CS62 cells.**

Me*BRCA1* patterns were assessed in **A.** WEHI-CS62 and **B.** post-DNMT1i treated cell population or **C-F.** individual FACS sorted clones using targeted bisulfite next generation sequencing (meNGS) of the *BRCA1* promoter. Individual epialleles present at ≥5% frequency in the sample are presented, along with a merged summary of less frequent epialleles (<5%). Each circle represents CpG sites along the *BRCA1* promoter, and numbers on x-axis indicate CpG distance from *BRCA1* transcription start site.

**Figure S6**

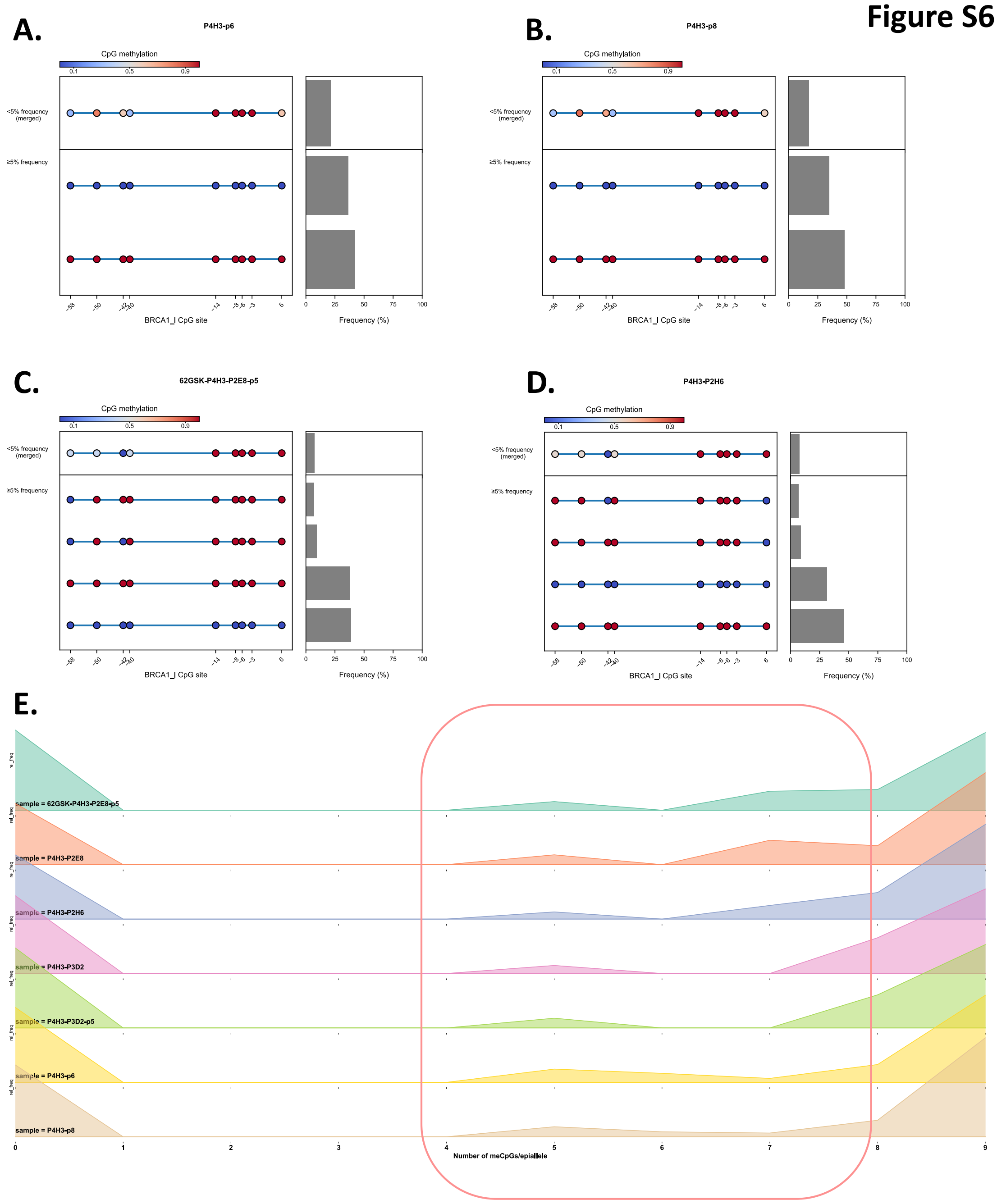


**Supplementary Figure S6. Heterogenous *BRCA1* methylation loss retained in second round of clones generated from post-DNMT1i WEHI-CS62 clone P4H3.**
Me*BRCA1* patterns were assessed in **A.** Passage p6 and **B.** p8 of post-DNMT1i WEHI-CS62 clone P4H3 using targeted bisulfite next generation sequencing (meNGS) of the *BRCA1* promoter. P4H3 appeared to have similar frequencies of fully methylated and fully unmethylated epialleles. A single cell FACS sort of this clonal population after 4 months in culture resulted in new clones with similar methylation patterns, with example meNGS of **C.** P2E8 and **D.** P2H6 shown here. Individual epialleles present at ≥5% frequency in the sample are presented, along with a merged summary of less frequent epialleles (<5%). Each circle represents CpG sites along the *BRCA1* promoter, and numbers on x-axis indicate CpG distance from *BRCA1* transcription start site. **E.** Analysis of *BRCA1* meNGS from these clones, and clone P3D2 from the sort of P4H3, revealed a mostly bi-modal distribution of epialleles, but with some low-frequency partially-methylated epialleles (in red box) present in all samples.

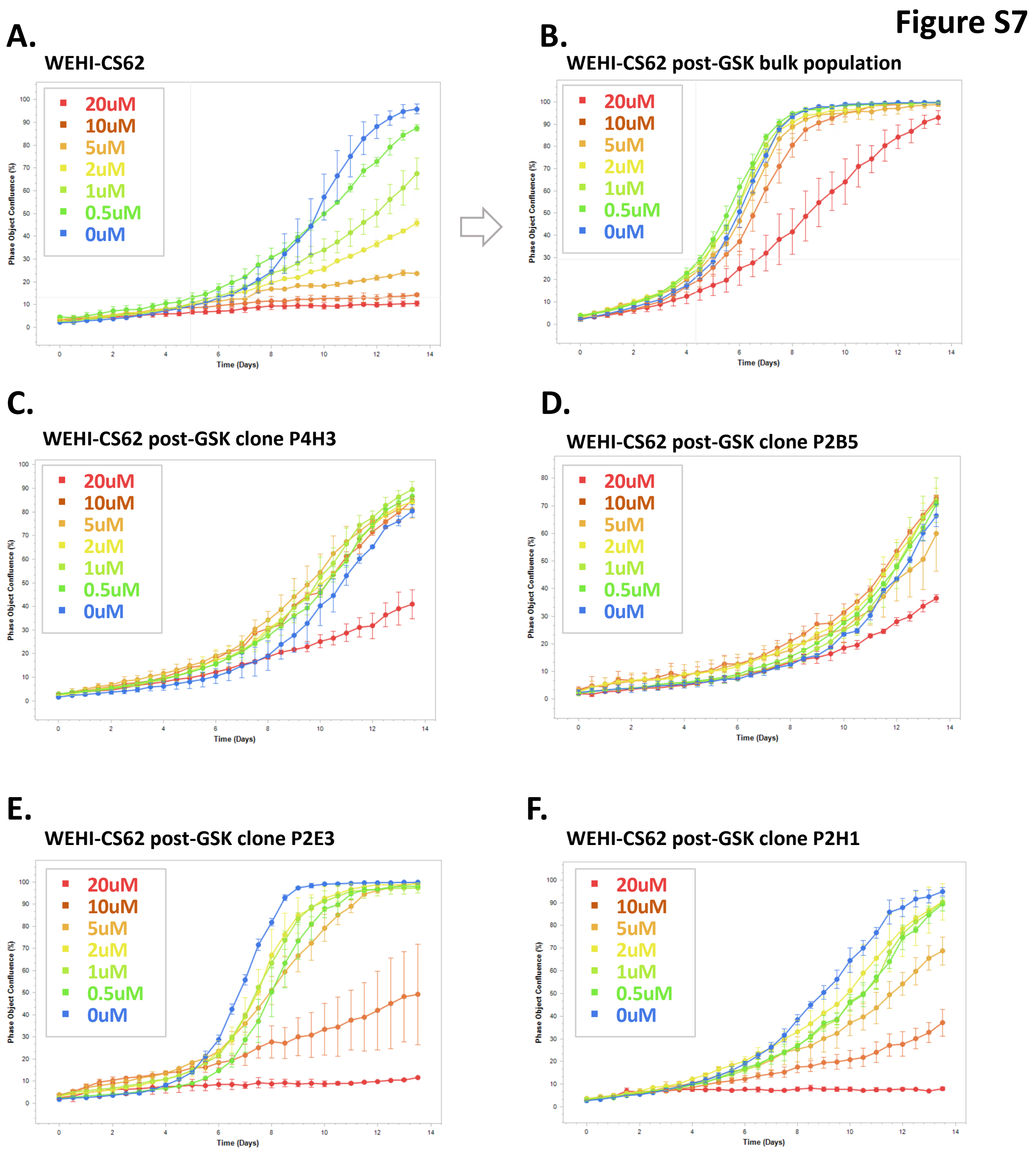


**Supplementary Figure S7. PARPi rucaparib responses of clones sorted from DNMT1i-treated WEHI-CS62.**

The **A.** parental WEHI-CS62 cells, **B.** post-DNMT1i bulk population and **C-F.** various clonal lines derived from FACS of the post-DNMT1i bulk population were treated with the indicated doses of PARPi and confluence was measured over 14 days on the Incucyte live cell imaging platform (cells imaged every 6 hours). Each point is a mean value of two technical replicates +/- SEM. Experimental replicates (n=3) presented in Supplementary Figure S8.

#### Figure S8

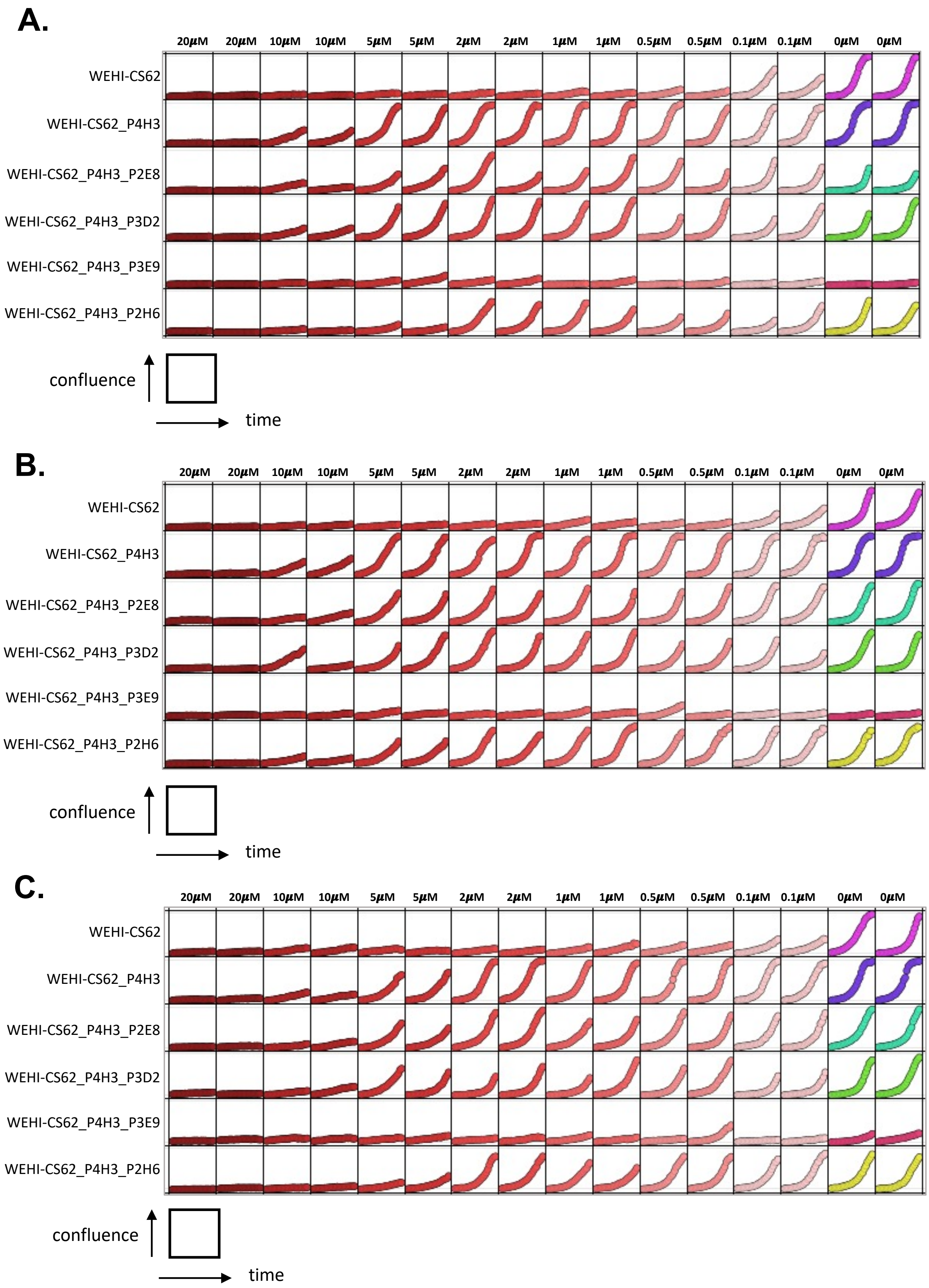


**Supplementary Figure S8. Experimental replicates of PARPi rucaparib responses of clones sorted from DNMT1i-treated WEHI-CS62.**

The parental WEHI-CS62 cells, post-DNMT1i bulk population, post-DNMT1i clone P4H3 from first FACS sort and various clonal lines derived from the second FACS sort (of P4H3) were treated with the indicated doses of PARPi and confluence was measured over 14 days on the Incucyte live cell imaging platform (cells imaged every 6 hours). Experimental replicates 1-3 are presented in panels **A.** to **C.,** respectively. Each box represents an imaged well on a plate. The confluence is measured on the y-axis, and time on the x-axis.

### Figure S9

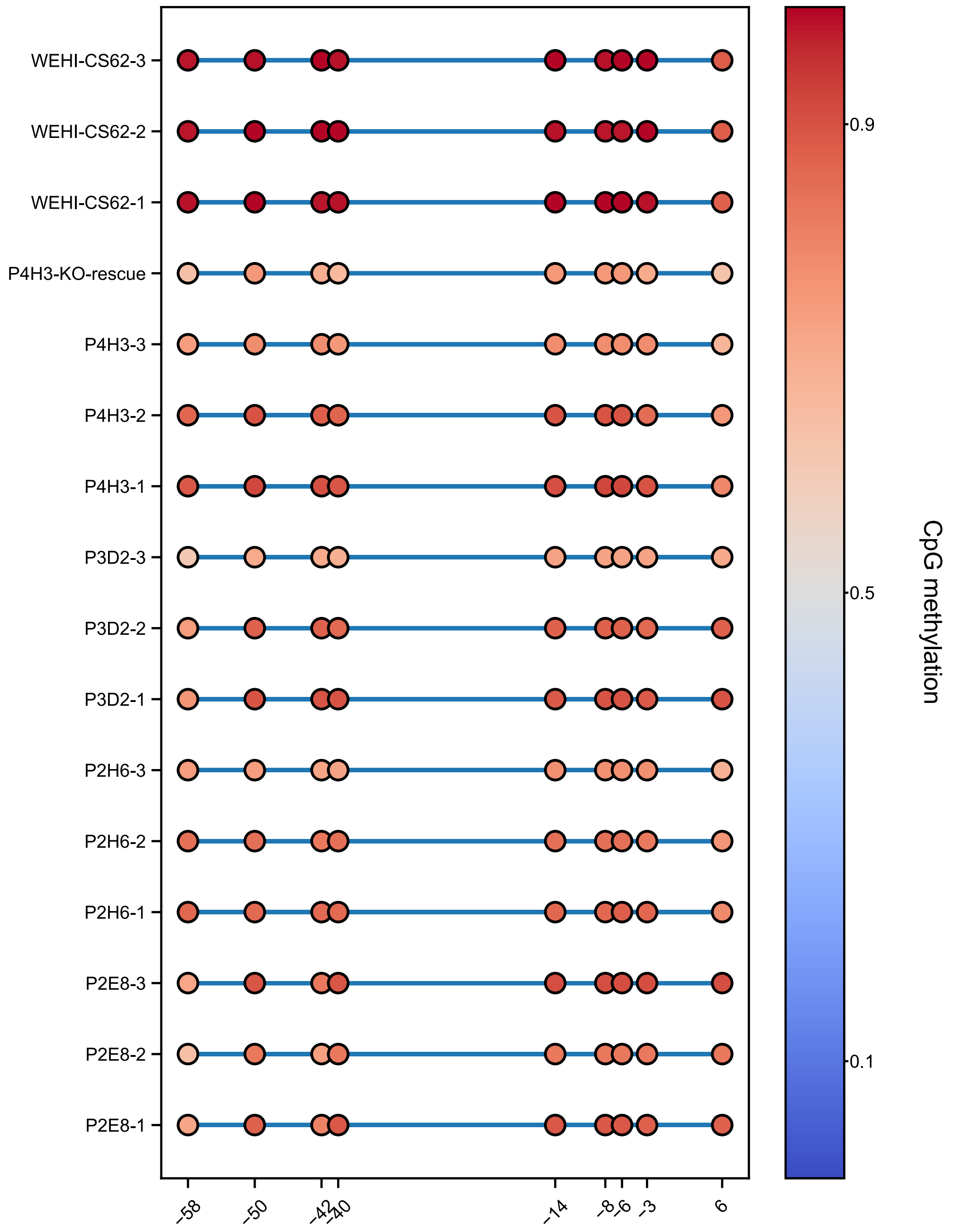


**Supplementary Figure S9. Targeted meNGS of *BRCA1* promoter in clones sorted from DNMT1i-treated WEHI-CS62.**

The *BRCA1* sequencing methylation results correspond to the samples presented in Supplementary Figure S8. Cell line and clone names are indicated, and the suffix or _1, _2 or _3 refers to the experimental replicate (for n=3 total). Each circle represents average methylation level for a CpG site along the *BRCA1* promoter, and numbers on x-axis indicate CpG distance from *BRCA1* transcription start site. The colour represents the average degree of methylation at each site.

#### Figure S10

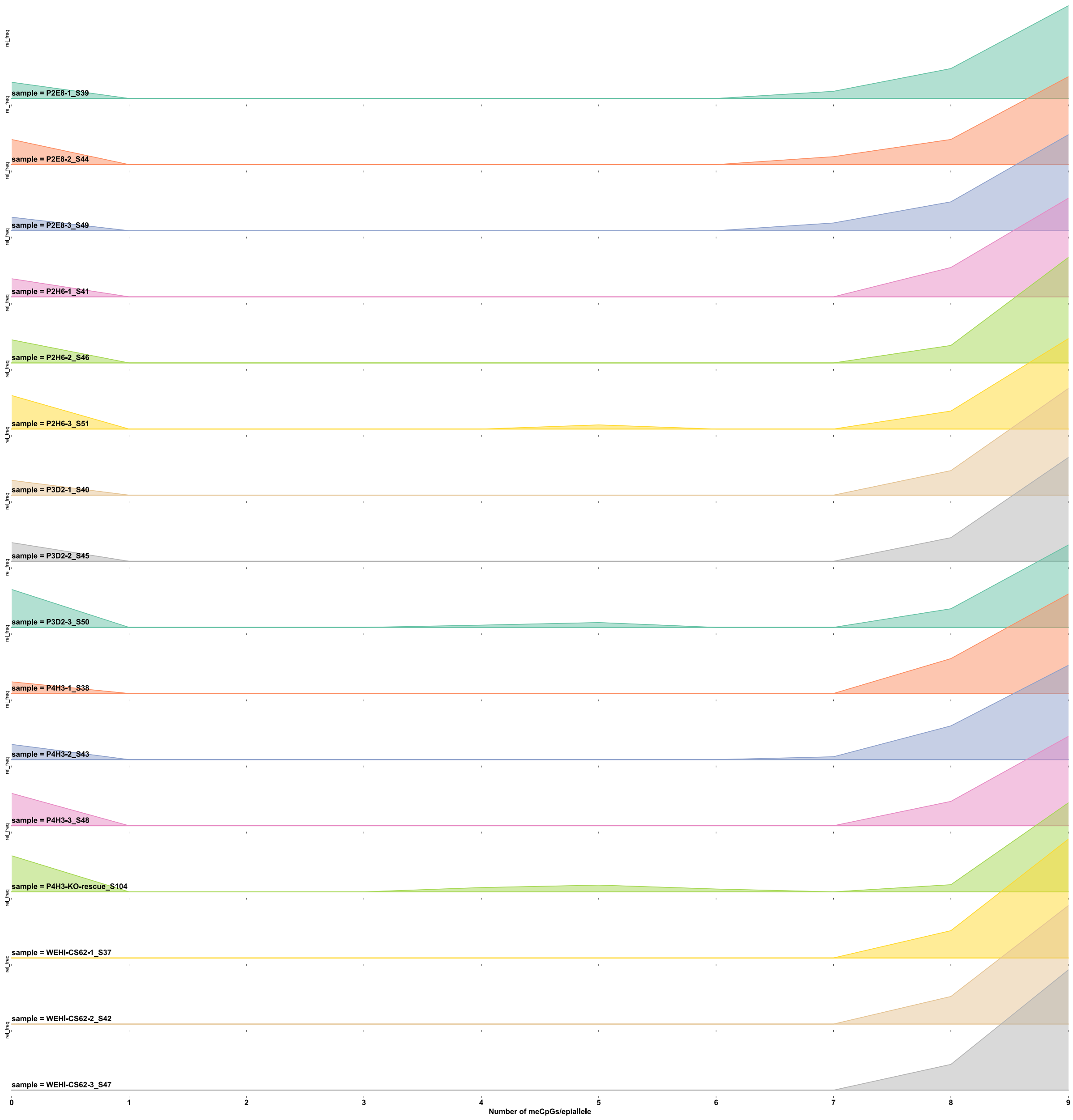


**Supplementary Figure S10. Targeted meNGS of *BRCA1* promoter in clones sorted from DNMT1i-treated WEHI-CS62 presented as epiallele distribution.**

The *BRCA1* sequencing methylation results correspond to the samples presented in Supplementary Figure S8. Cell line and clone names are indicated, and the suffix or -1, -2 or -3 refers to the experimental replicate (for n=3 total), with number following underscore indicating the meNGS code (from multiplexing). Y-axis = frequency of each me*BRCA1* epiallele, numbers on x-axis indicate the number of methylated CpG sites per epiallele (0 = fully unmethylated and 9= fully methylated).

### Figure S11

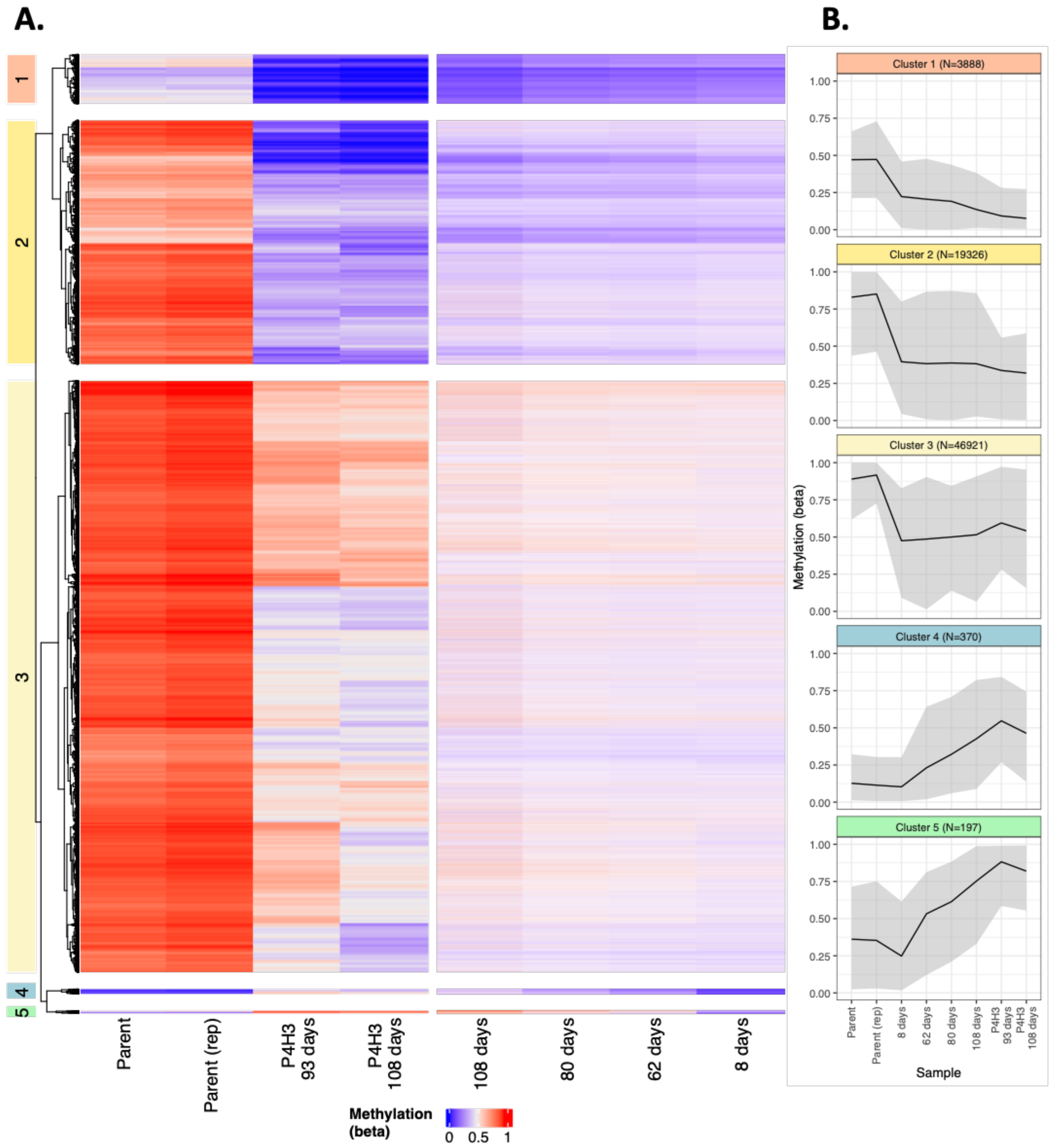


**Supplementary Figure S11. Long-term global methylation changes in post-DNMT1i WEHI-CS62 cells.**

Differentially methylated (DM) probes (70,702/828,647, $\Delta\beta \geq 0.25$) were identified by comparing parental WEHI-CS62 cells with clonal P4H3 and long-term recovered bulk samples (108 days post-treatment). **A.** Distinct methylation patterns were revealed through hierarchical clustering of untreated parental and P4H3 clones, showing five DM groups. The changes in the corresponding probes over eight to 108 days of recovery are depicted in the right panel of the heatmap. **B.** Methylation trends over time were tracked and summarized as median beta values (black line), with maximum and minimum variations shaded in grey. Clusters 1, 2 and 3 showed decreased methylation compared to parental line, while clusters 4 and 5 showed increased methylation, all stabilizing in P4H3.

**Figure S12**

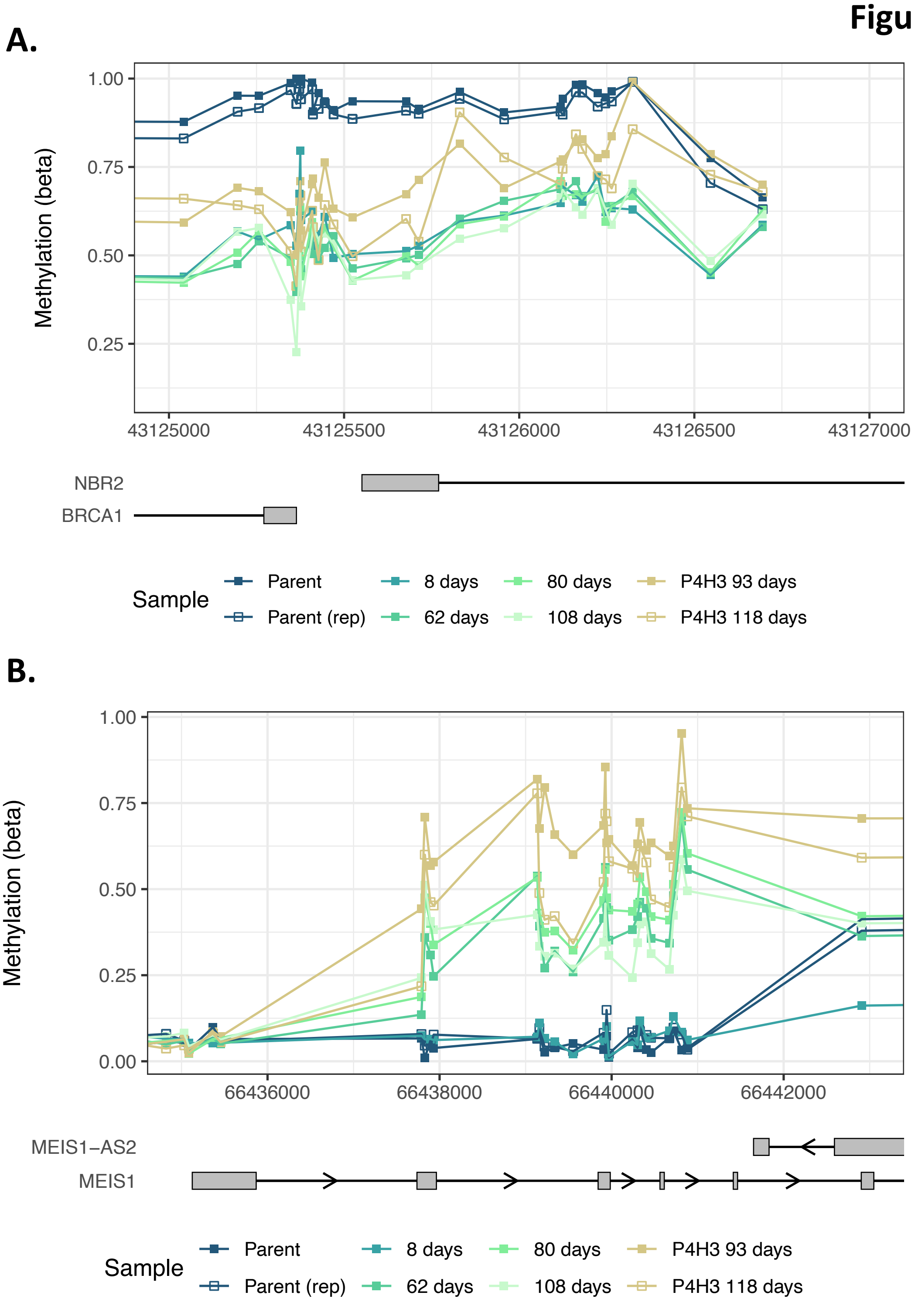


**Supplementary Figure S12. Methylation of *BRCA1* and an example gene *MEIS1* with increased methylation post-DNMT1i data.**

**A.** CpG probe methylation values across the *BRCA1* promoter region (local genes and chromosomal locations indicated below) for the various post-DNMT1i timepoints of WEHI-CS62 (including clone P4H3). **B.** Unlike the *BRCA1* promoter, the *MEIS1* gene region had an increase in methylation following DNMT1i treatment of WEHI-CS62 (cluster 4 and 5 from differential methylation analysis).

**Figure S13**

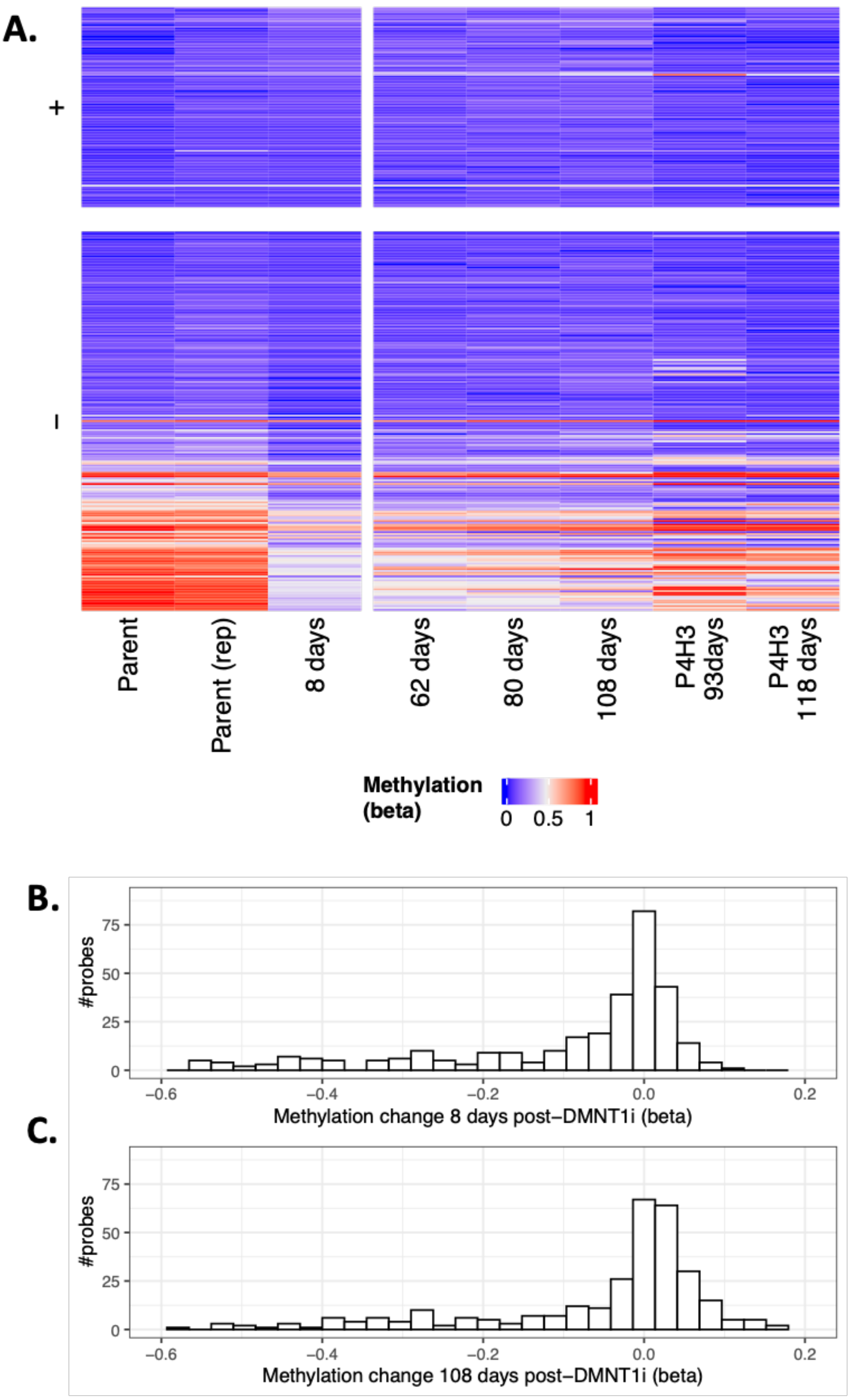


**Supplementary Figure S13. Methylation status of CpG sites reported in Giri et al. 2019 with increased methylation following decitabine treatment.**

**A.** Of the 638 CpGs previously reported by Giri & Aittokallio [2] to have increased methylation five days post-decitabine treatment, 313 matched probes were available in WEHI-CS62. Eight days post-DNMT1i, a subset (34%, 108/313) exhibited a slight methylation increase ($\Delta\beta < 0.2$, labeled “+”), while the majority (66%, 205/313) showed methylation loss (labeled “–”). **B.** Distribution of methylation changes in the eight-day post-DNMT1i samples compared to untreated controls further emphasizes the shift towards methylation loss, in

contrast to the methylation increases reported by Giri & Aittokallio.
**C.** Minimal increases in methylation were observed even with extended recovery times (108 days). For B-C beta probe methylation change values are presented on the x-axis, with 0 representing no change.

### Figure S14

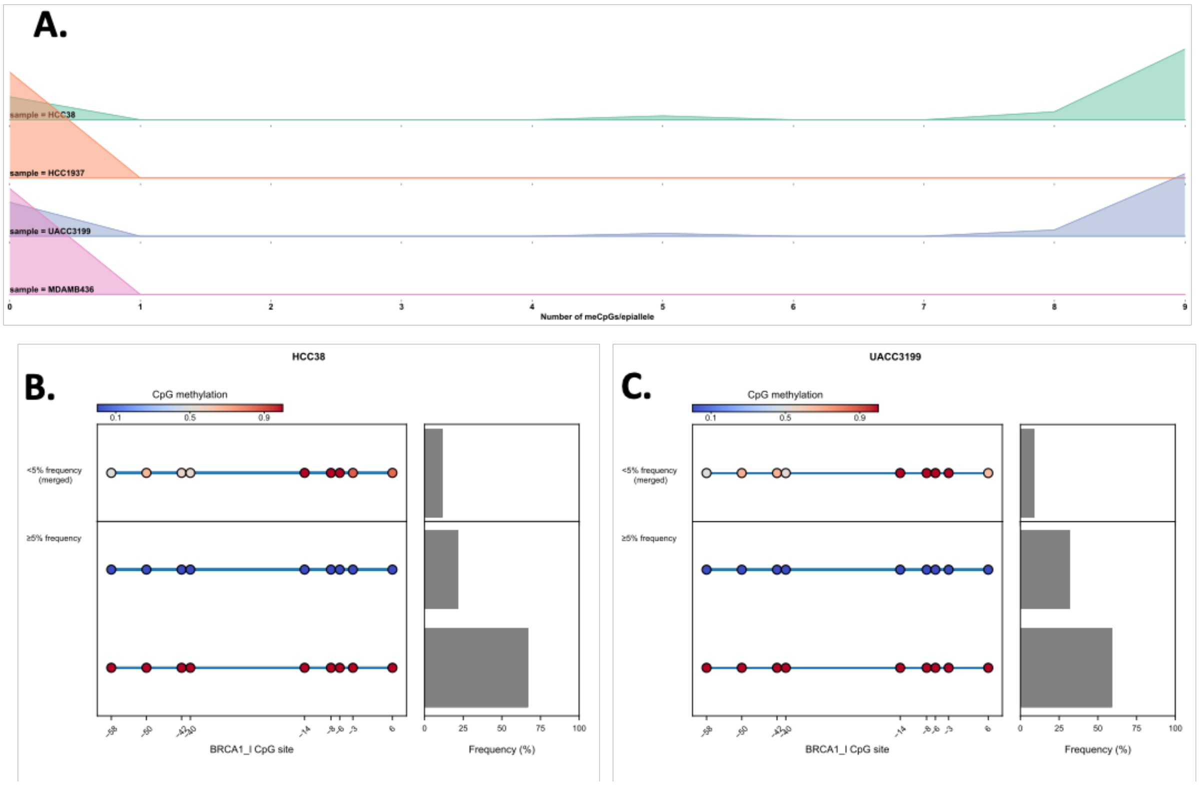


**Supplementary Figure S14. Breast cancer cell lines with heterozygous *BRCA1* methylation.**

**A.** *BRCA1* meNGS was performed for two breast cancer cell lines, HCC38 and UACC3199, previously reported to have me*BRCA1*, and two control cell lines lacking me*BRCA1*, HCC1937 and MDA-MB-436. Unmethylated epialleles were detected in both HCC38 and UACC3199 suggesting heterozygous *BRCA1* methylation. This is also illustrated by individual sample meNGS plots for **B.** HCC38 and **C.** UACC3199. Individual epialleles present at ≥5% frequency in the sample are presented, along with a merged summary of less frequent epialleles (<5%). Each circle represents CpG sites along the *BRCA1* promoter, and numbers on x-axis indicate CpG distance from *BRCA1* transcription start site.

#### Figure S15

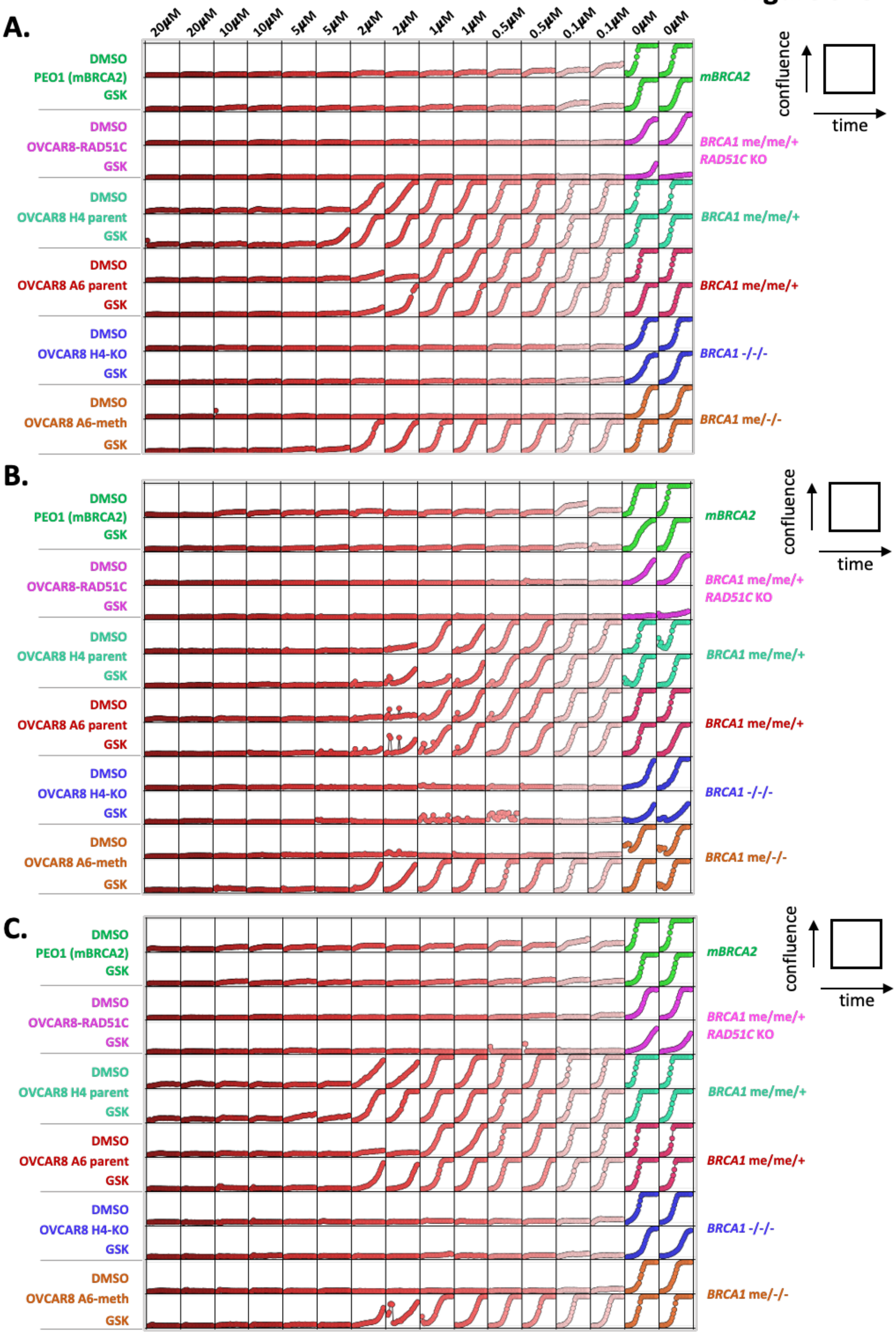
A.
20μM
20μM
10μM
10μM
5μM
5μM
2μM
2μM
1μM
1μM
0.5μM
0.5μM
0.1μM
0.1μM
0μM
0μM
DMSO
PEO1 (mBRCA2)
GSK
DMSO
OVCAR8-RAD51C
GSK
DMSO
OVCAR8 H4 parent
GSK
DMSO
OVCAR8 A6 parent
GSK
DMSO
OVCAR8 H4-KO
GSK
DMSO
OVCAR8 A6-meth
GSK
mBRCA2
BRCA1 me/me/+
RAD51C KO
BRCA1 me/me/+
BRCA1 me/me/+
BRCA1 -/-/-
BRCA1 me/-/-
confluence
time
B.
C.

**Supplementary Figure S15. Experimental replicates of PARPi rucaparib responses of various DNMT1i-treated cell lines.**

Various cell lines (including homozygous me*BRCA1* OVCAR8 A6-meth) were treated with the indicated doses of PARPi rucaparib and confluence was measured over 14 days on the Incucyte live cell imaging platform (cells imaged every 6 hours). Experimental replicates 1-3 are presented in parts **A. to C.,** respectively. Each box represents an imaged well on a plate, and the confluence is measured on the y-axis, and time on the x-axis, as indicated by the boxes to the left of each section. The OVCAR8-RAD51C cell line (with knockout of *RAD51C* [3]), did not grow well following DNMT1i GSK3484862 (GSK) compared to DMSO control. PEO1 has a mutation in *BRCA2* (m*BRCA2*) and is sensitive to PARPi.

#### Figure S16

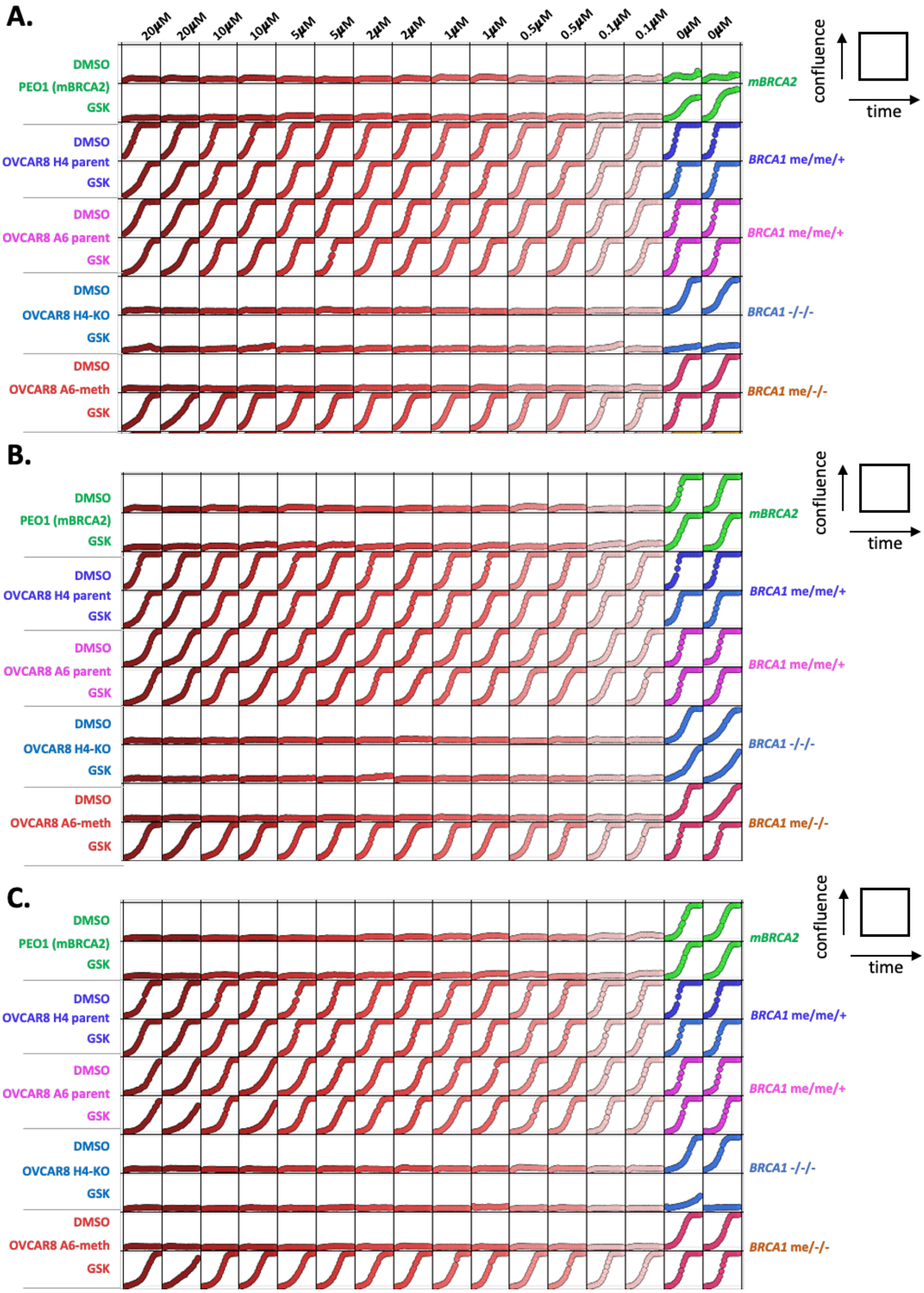
A.
20μM
20μM
10μM
10μM
5μM
5μM
2μM
2μM
1μM
1μM
0.5μM
0.5μM
0.1μM
0.1μM
0μM
0μM
DMSO
PEO1 (mBRCA2)
GSK
mBRCA2
DMSO
OVCAR8 H4 parent
GSK
BRCA1 me/me/+
DMSO
OVCAR8 A6 parent
GSK
BRCA1 me/me/+
DMSO
OVCAR8 H4-KO
GSK
BRCA1 -/-/-
DMSO
OVCAR8 A6-meth
GSK
BRCA1 me/-/-
confluence
time
B.
DMSO
PEO1 (mBRCA2)
GSK
mBRCA2
DMSO
OVCAR8 H4 parent
GSK
BRCA1 me/me/+
DMSO
OVCAR8 A6 parent
GSK
BRCA1 me/me/+
DMSO
OVCAR8 H4-KO
GSK
BRCA1 -/-/-
DMSO
OVCAR8 A6-meth
GSK
BRCA1 me/-/-
confluence
time
C.
DMSO
PEO1 (mBRCA2)
GSK
mBRCA2
DMSO
OVCAR8 H4 parent
GSK
BRCA1 me/me/+
DMSO
OVCAR8 A6 parent
GSK
BRCA1 me/me/+
DMSO
OVCAR8 H4-KO
GSK
BRCA1 -/-/-
DMSO
OVCAR8 A6-meth
GSK
BRCA1 me/-/-
confluence
time

**Supplementary Figure S16. Experimental replicates of potent PARPi AZD5305 responses of various DNMT1i-treated cell lines.**

Various cell lines (including homozygous me*BRCA1* OVCAR8 A6-meth) were treated with the indicated doses of potent PARPi AZD5305 and confluence was measured over 14 days on the Incucyte live cell imaging platform (cells imaged every 6 hours). Experimental replicates 1-3 are presented in parts **A. to C.,** respectively. Each box represents an imaged well on a plate, and the confluence is measured on the y-axis, and time on the x-axis, as indicated by the boxes to the left of each section. The OVCAR8 H4-KO cell line (with full knockout of *BRCA1* [4]) did not grow well following DNMT1i GSK3484862 (GSK) compared to DMSO control in two replicates. PEO1 has a mutation in *BRCA2* (m*BRCA2*) and is sensitive to PARPi.

Figure S17

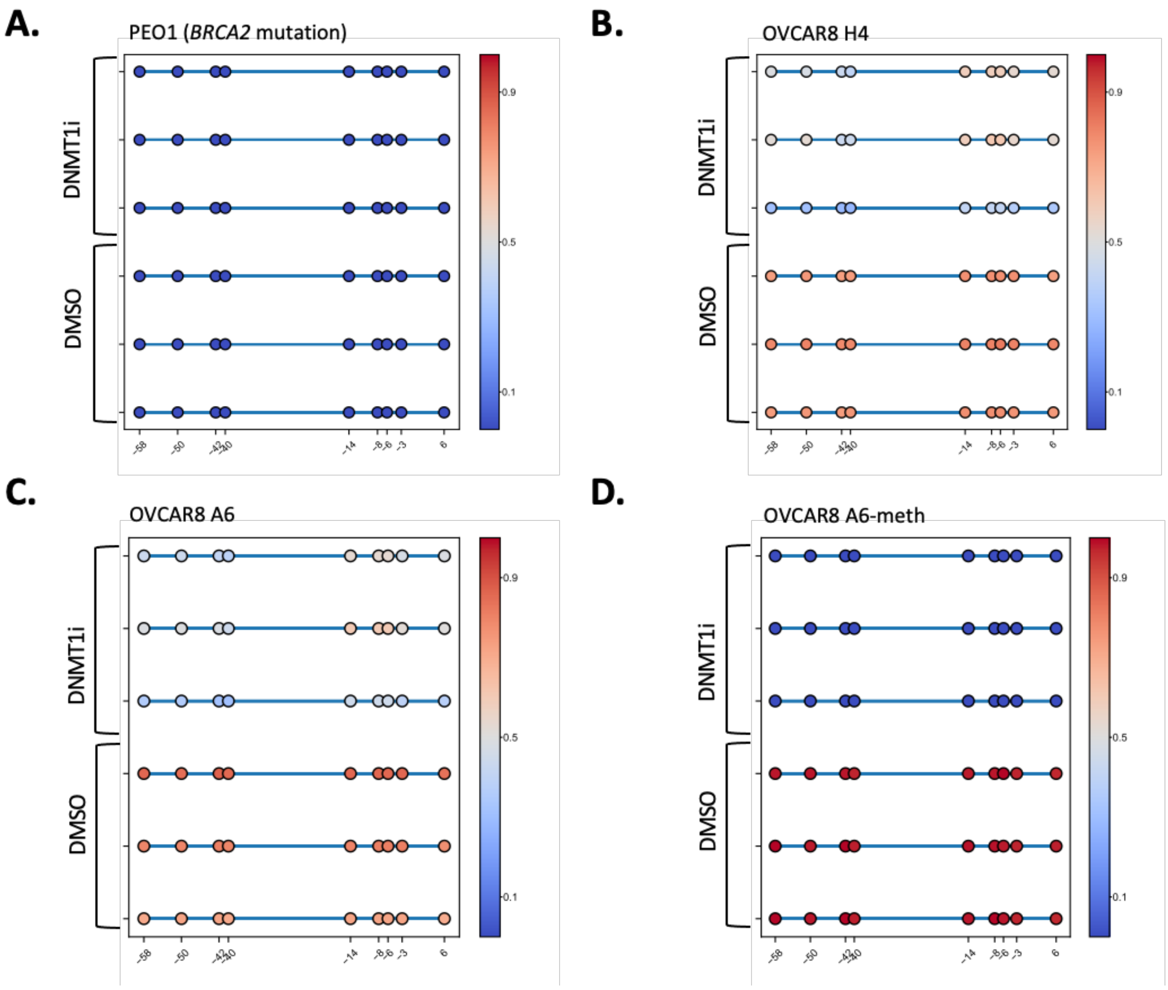


**Supplementary Figure S17. Methylation of *BRCA1* in various DNMT1i-treated cell lines.** These *BRCA1* sequencing methylation results correspond to three replicates (passages) of the cell lines presented in Supplementary Figures S14-15, treated with either DNMT1i GSK3484862 or DMSO control. **A.** Results for cell line PEO1 (m*BRCA2* = *BRCA2* mutation). **B.** Results for cell line OVCAR8 H4 (landing pad parental line used to create *BRCA1* KO line OVCAR8 H4-KO). **C.** Results for cell line OVCAR8 A6 (landing pad parental line used to create homozygous *BRCA1* methylation line OVCAR8 A6-meth). **D.** Results for cell line OVCAR8 A6 OVCAR8 A6-meth. Each circle represents CpG sites along the *BRCA1* promoter, and numbers on x-axis indicate CpG distance from *BRCA1* transcription start site. The colour represents the average degree of methylation at each site.

### Figure S18

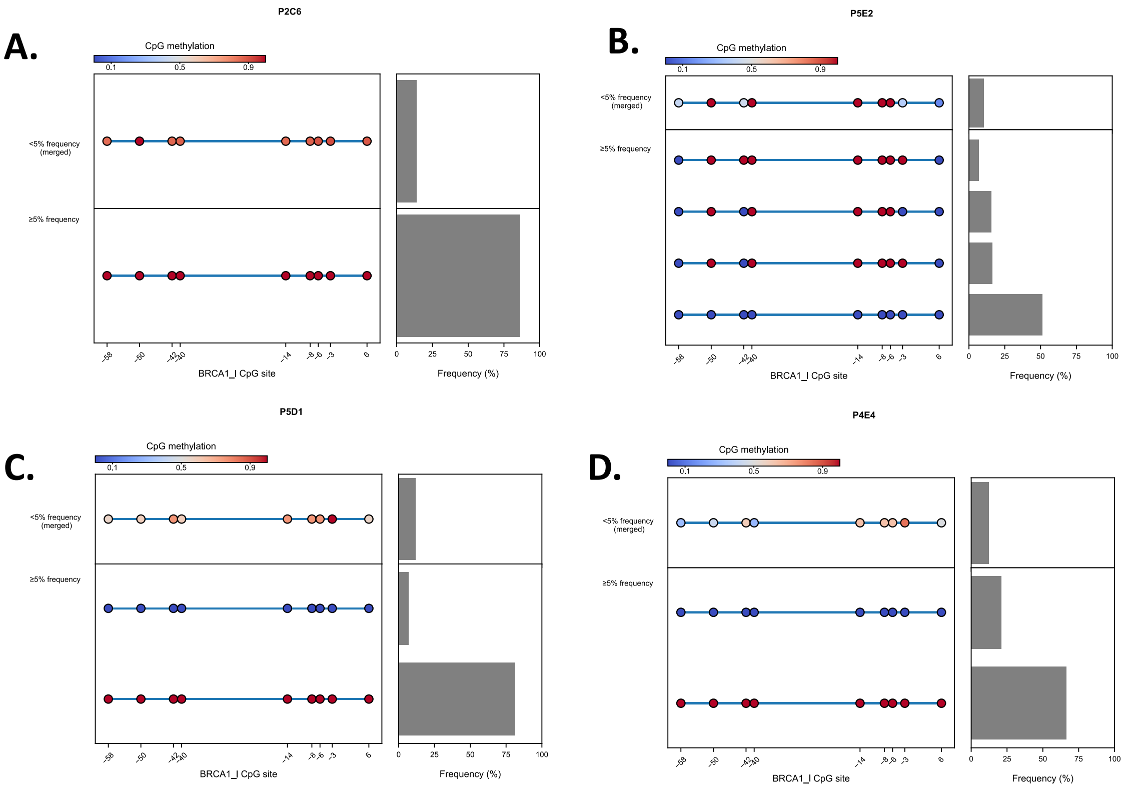


**Supplementary Figure S18. *BRCA1* methylation in clones generated following CRISPR knock-out of *BRCA1* gene in post-DNMT1i WEHI-CS62 clone P4H3.**
Targeted CRISPR deletion of the full *BRCA1* gene using dual guide RNAs targeting either side of the *BRCA1* gene (encompassing the promoter) was carried out in clone P4H3 (~50% methylation) derived from DNMT1i treated WEHI-CS62. No homozygous KO clones were identified. **A.** One clone (P2C6) was found to have homozygous *BRCA1* methylation by targeted methylation NGS. **B-C.** Other knock-out clones screened were found to retain unmethylated epialleles of *BRCA1* and were not investigated further. Individual epialleles present at ≥5% frequency in the sample are presented, along with a merged summary of less frequent epialleles (<5%). Each circle represents CpG sites along the *BRCA1* promoter, and numbers on x-axis indicate CpG distance from *BRCA1* transcription start site.

#### Figure S19

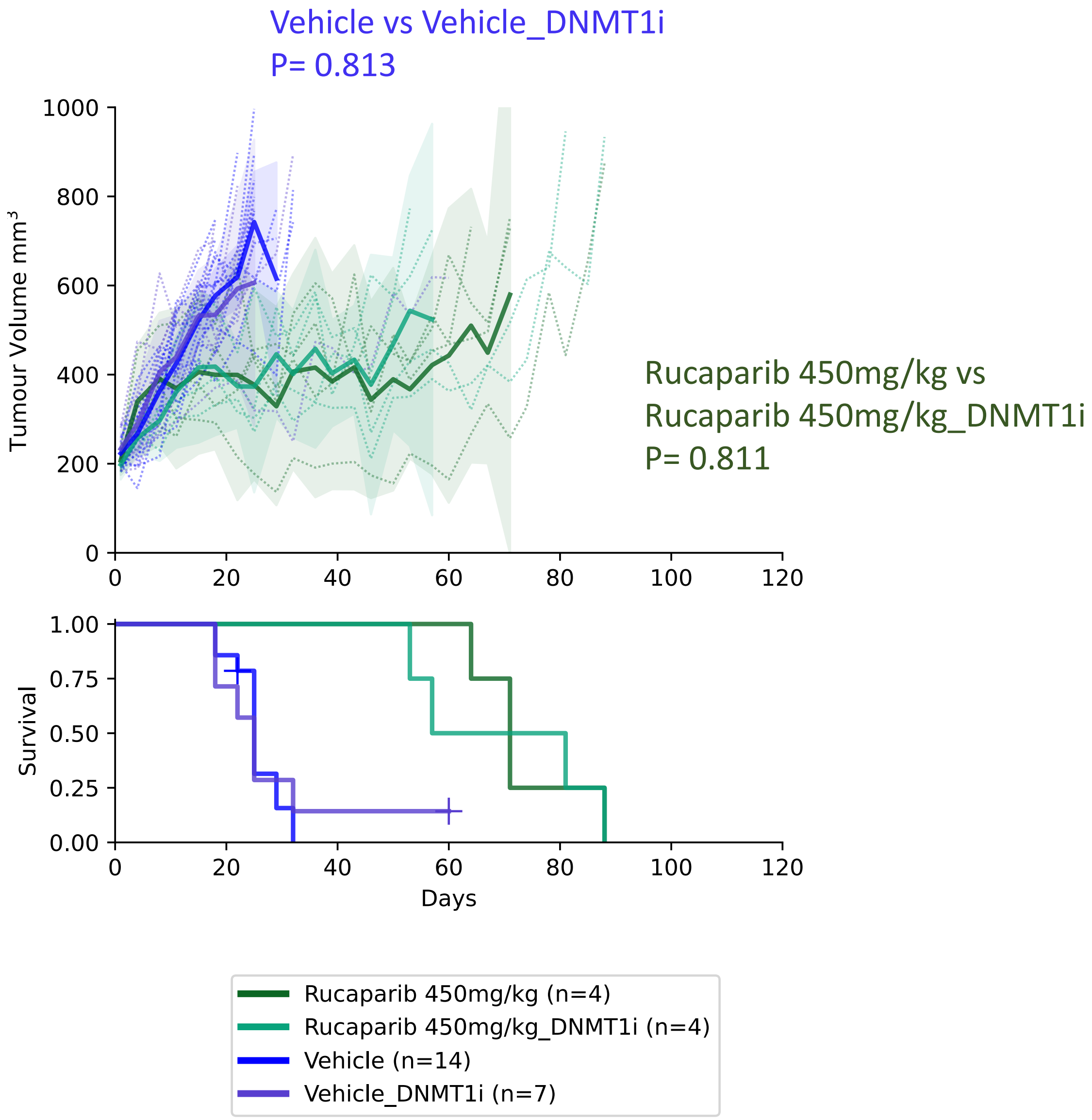


**Supplementary Figure S19. Comparison of PARPi rucaparib responses in standard PDX #62 lineage and the post-DNMT1i lineage.**

No significant difference in vehicle or PARPi tumour growth rates was observed following *in vivo* DNMT1i (GSK3685032) treatment of PDX #62. Mean PDX tumour volume (mm3) ± 95% CI (hashed lines are individual mice) and corresponding Kaplan–Meier survival analysis. Censored events are represented by crosses on Kaplan–Meier plot; n = individual mice. Log-Rank test P values for each comparison are presented.

### Figure S20

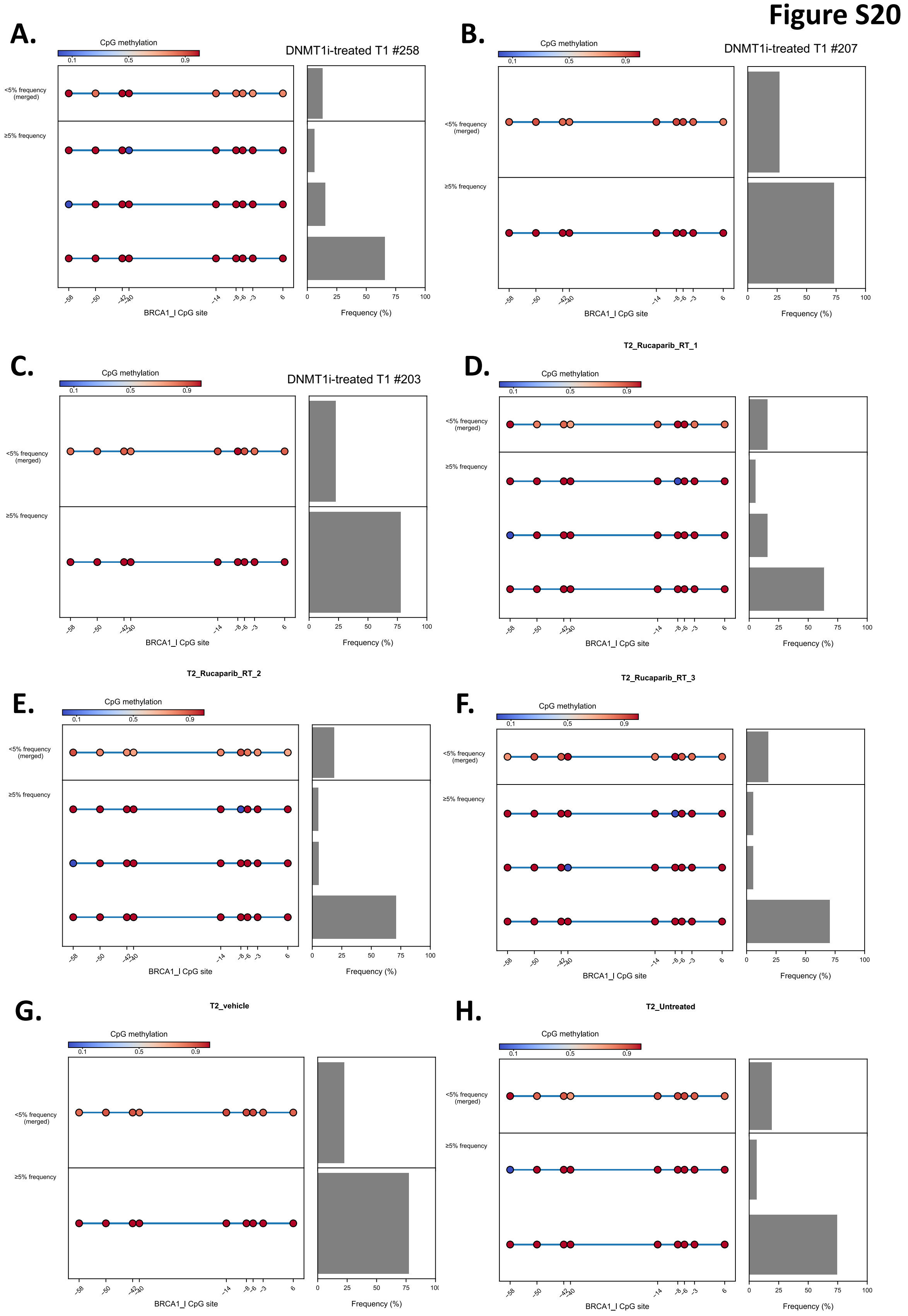


**Supplementary Figure S20.** ***BRCA1*** **methylation results for PDX #62 tumours following DNMT1i** ***in vivo*** **treatment.**

**A-C.** No unmethylated *BRCA1* epialleles were observed in the three PDX #62 tumours treated with DNMT1i GSK3685032 *in vivo*. **D-F.** No unmethylated *BRCA1* epialleles were observed in two of the three PARPi rucaparib treated PDX #62 tumours from the post-DNMT1i (GSK3685032) PDX lineage, with one tumour showing 2% unmethylated epialleles (**E**). **G.** No methylation loss was observed in one of two untreated PDX #62 tumours from the post-DNMT1i (GSK3685032) PDX lineage, or in **H.** the vehicle-treated tumour from this lineage. Individual epialleles present at ≥5% frequency in the sample are presented, along with a merged summary of less frequent epialleles (<5%). Each circle represents CpG sites along the *BRCA1* promoter, and numbers on x-axis indicate CpG distance from *BRCA1* transcription start site. T1 indicates PDX transplant/passage 1, and T2 indicates PDX transplant/passage 2.
